# Dynamic Lake Ice Conditions Shape Caribou Water-Crossing Behavior in the Arctic

**DOI:** 10.1101/2025.09.30.678981

**Authors:** Qianru Liao, Eliezer Gurarie, William F. Fagan

## Abstract

Successful animal migration hinges on navigation and decision-making in dynamic environments. Yet, how individuals navigate transient, fine-scale landscape barriers, such as seasonally ice-covered water bodies, remains poorly understood. Understanding these responses is critical for forecasting migration routes and connectivity under global change. In the Arctic, rising temperatures are causing earlier ice melt and later freeze-up, reshaping landscape permeability and potentially disrupting migration routes for overland migrants, such as barren-ground caribou *(Rangifer tarandus)*, a keystone Arctic species, which relies on frozen lakes and rivers for efficient spring travel to calving grounds. While caribou generally prefer ice to open water, behavioral responses to changing ice conditions have not been quantitatively assessed. We analyzed 20 years (2001-2021) of GPS data for 406 adult caribou and daily MODIS land surface albedo to examine lake-crossing decisions at Contwoyto Lake, a long (>100 km) glacial lake in northern Canada. We classified transit events as crossing or circumnavigation based on GPS trajectories relative to lake boundaries and linked behavioral decisions to spatially and temporally resolved ice conditions. Our models revealed distinct seasonal drivers. Spring crossing decisions were shaped by intermediate-scale ice conditions, with a behavioral threshold at a path-averaged annual albedo percentile rank of 0.56, corresponding to intermediate late-spring melt conditions when lake ice transitions from continuous cover toward fragmented surfaces. In fall, when the lake was ice-free, movement-related factors such as relative speeds along alternative routes better explained behavior. Our findings show how ice acts as a seasonal behavior filter, shaping functional connectivity through perceptual and energetic constraints. Although developed for caribou, this framework is transferable across species and systems. By linking high-resolution, spatiotemporal remote sensing to individual behavior, our framework identifies quantitative behavioral thresholds in response to dynamic, climate-sensitive landscape features, supporting predictive monitoring of climate-driven shifts in migratory behavior and emerging constraints on movement.

**Open Research Statement:** The MODIS datasets used in this study are publicly available from NASA’s Land Processes Distributed Active Archive Center (LP DAAC). Daily shortwave black-sky albedo values were obtained from MCD43A3.061, and land cover types from MCD12Q1.061. Caribou GPS collar data were provided by the Government of the Northwest Territories, Department of Environment and Climate Change (GNWT-ECC), and are not publicly available due to data-sharing restrictions. Qualified researchers may request access from GNWT-ECC (https://www.gov.nt.ca/ecc/en). The code used for data processing and analysis will be archived in GitHub upon manuscript acceptance.

## 1. Introduction

Seasonal long-distance migration is a fundamental ecological strategy that enables animals to track dynamic resources, avoid predators, and access critical reproductive habitats (Fryxell and Sinclair 1988b; Avgar et al. 2014). This strategy allows larger populations to live and reproduce in regions that are only seasonally suitable (Fryxell and Sinclair 1988a; 1988b; Fryxell et al. 1988). However, climate change and anthropogenic development are altering the spatial and temporal distribution of resources, challenging the persistence of migratory species that depend on predictable large-scale movements (Wilcove and Wikelski 2008; Harris et al. 2009; Chen et al. 2011; Middleton et al. 2013; Xu et al. 2021; Kauffman et al. 2021; Sutherland 1998).

While some adaptations of migratory species to changing environmental conditions may be genetically based (Alerstam et al. 2003; Anderson et al. 2013), long-lived social migrants often rely more heavily on a combination of social interactions —from older members to newer ones—and shared memories to navigate new settings (Mueller et al. 2013; Teitelbaum et al. 2016; Berdahl et al. 2018; Jesmer et al. 2018; Gurarie et al. 2021; Aikens et al. 2022). For example, recent work documented a 500-km winter range shift in the Western Arctic caribou herd, apparently shaped by the collective memory of poor conditions in previously favored areas (Gurarie et al. 2024). While this behavioral flexibility reflects the potential for adaptive reconfiguration of movement, it also reveals imperfect outcomes, highlighting how even socially transmitted strategies may struggle to keep pace with rapid environmental change. Despite these insights, important knowledge gaps remain. While a growing body of work has demonstrated that migratory ungulates track spatiotemporal variation in forage resources during migration (Merkle et al. 2016; Aikens et al. 2017, 2020; Middleton et al. 2018; Abrahms et al. 2021; Kauffman et al. 2021; Laforge et al. 2021), comparatively little attention has been paid to how migrants navigate dynamic physical barriers, such as ice-covered water bodies, whose permeability shifts rapidly with seasonal and climatic conditions. Understanding these tactical movement decisions in response to transient landscape features is critical for forecasting connectivity under climate change, particularly in Arctic systems where snow and ice, rather than vegetation phenology, govern movement during much of the year (Boelman et al. 2019; Matias et al. 2024).

In cold regions, seasonal ice serves as a key environmental feature that modulates animal movement. Acting as a temporary bridge across aquatic barriers, ice forms vital migration corridors for terrestrial species (Banfield 1954b). However, with Arctic warming proceeding nearly four times faster than the global average (Screen and Simmonds 2010; Jeong et al. 2014; Rantanen et al. 2022), the timing and reliability of ice formation have become increasingly erratic. Long-term observational records across the Northern Hemisphere reveal earlier break-up and later freeze-up of lakes and rivers (Magnuson et al. 2000), shortening the seasonal window for ice-supported movement. These shifting ice regimes threaten the functional connectivity of aquatic-terrestrial landscapes (Leblond et al. 2016), increasing detour costs, delaying access to calving grounds, and elevating mortality risks during crossings (Miller and Gunn 1986).

Caribou and reindeer (both *Rangifer tarandus*), widespread circumpolar ungulates (*family Cervidae*) in Arctic areas (Feldhamer et al. 2003; Hummel and Ray 2008), exemplify the challenges posed by changing ice conditions. These migratory ungulates undertake long-distance terrestrial migrations of up to 1000 km annually between calving and wintering grounds (Fancy et al. 1989; Berger 2004; Joly et al. 2019), often relying on ice-covered water bodies as critical overland migration pathways. Some populations, such as the Peary caribou (Jenkins et al. 2016; Gautier et al. 2022) and the Dolphin and Union caribou (Poole et al. 2010), even migrate across sea ice between islands, underscoring the ecological importance of ice as connective corridors.

Although caribou are capable swimmers, open water, particularly during ice breakup, presents a more hazardous and energetically costly environment for movement. For example, caribou that break through poor-quality ice may suffer severe hemorrhage or trauma when attempting to escape or even succumb to shock and exposure (Miller and Gunn 1986). Even after escaping, weakened individuals are more vulnerable to predation (Banfield 1954a). In contrast, stable ice surfaces provide flat, vegetation-free travel routes that facilitate rapid movement and predator detection (Mysterud and Østbye 1999). As ice loss accelerates, open water bodies increasingly act as dispersal barriers, forcing caribou to either make long detours (Leblond et al. 2016) or swim, which is less efficient and increases the risk of exhaustion (Fish 1994). Energy loss and delays are especially consequential during spring migration, when delayed arrivals may disrupt highly synchronized birthing periods, reducing calf survival and affecting population dynamics (Gurarie et al. 2019; Couriot et al. 2023). In contrast, fall movements tend to be more nomadic and flexible. The dramatic population decline of the Bathurst herd, from approximately 480,000 individuals to fewer than 7,000 over the past three decades (Adamczewski et al. 2022), alongside broader global declines in ungulate migrations (Harris et al. 2009; Tucker et al. 2018), underscores the urgency of understanding how shifting ice phenology affects migratory behavior in the Arctic. Despite growing interest in Arctic lake ice trends (Livingstone 2000; Benson et al. 2012; Sharma and Magnuson 2014), most phenological studies rely on long-term in-situ records from large, well-monitored lakes. These observations typically lack consistent definitions for freeze-up and break-up dates (Eklund 1999; Magnuson et al. 2000; see Appendix S1), overlook within-lake spatial variation, limiting their relevance for understanding localized environmental cues. Yet such fine-scale variation may be critical for migratory animals making real-time movement decisions. To address these challenges, a variety of remote sensing approaches have been employed to monitor ice dynamics in lakes and rivers (see Appendix S1). Moderate Resolution Imaging Spectroradiometer (MODIS) products offer a favorable balance of spatial resolution (500 m), temporal frequency (daily), and long-term availability from both Terra and Aqua satellites (launched in 1999 and 2002, respectively), making them particularly useful for assessing lake ice conditions relevant to animal movement. However, many existing related applications of MODIS rely on manual interpretation of imagery (Pavelsky and Smith 2004; Cooley and Pavelsky 2016) or simplify spatially complex ice patterns into single metrics, such as average reflectance or centroid values (Šmejkalová et al. 2016), thereby obscuring the patchiness and progression of melting that animals likely perceive and respond to. Capturing these within-lake spatial heterogeneities is essential to link remotely sensed environmental change with animal tactical movement responses during migration.

Here we address these gaps by integrating GPS tracking of caribou with a pixel-level albedo analysis derived from MODIS daily surface reflectance data from both Terra and Aqua satellites. This approach uniquely combines fine spatial resolution with daily updates, allowing us to quantify dynamic ice conditions at the scale animals actually perceive. Focusing on the Contwoyto Lake region of Nunavut, Canada, we examine how daily shifts in lake surface reflectance, as a proxy for ice condition, influence caribou water-crossing behavior. Our results reveal seasonal behavioral thresholds tied to ice melt and demonstrate how Arctic warming may erode traditional migratory corridors. This study offers a quantitative link between climate-sensitive landscape dynamics and tactical movement decisions during migration.

## 2. Materials and Methods

### 2.1 Study Area

Contwoyto Lake (65° 39’ 0" N, 110° 43’ 0" W) is a large (around 957 km^!^) remote Arctic lake, near the border between Nunavut and the Northwest Territories, Canada (Pienitz et al. 1997). The lake is more than 110 km long but no wider than 10 km, with a northwest-to-southeast orientation. The region is characterized by a tundra climate, with long, harsh winters (often below -30°C) and short, cool summers (average 10°C) (Government of the Northwest Territories Department of Transportation et al. 1999). Annual precipitation is low (200–250 mm), with roughly half falling as snow (De Beers Canada Inc. 2002). The area is underlain by continuous permafrost and is located north of the continental treeline. Vegetation is dominated by sedges, dwarf shrubs, mosses, and lichens typical of Low Arctic tundra ecosystems (Downing et al. 2012). Caribou from the Bathurst, Beverly, and Bluenose East herds regularly migrate across or around the lake (Figure 1). In spring, the lake lies directly along the highly endangered Bathurst herd’s route between wintering areas to the southwest and calving grounds to the northeast (Gunn et al. 2001), making it a recurrent and functionally important decision point in seasonal migration. In the Tłįchǫ Dene language, the term Kokèti Ekwoǫ́ (“*migratory caribou of Contwoyto Lake*”) highlights the deep cultural and ecological ties between Indigenous peoples and the region (Tłįchǫ Research and Training Institute 2018).

**Figure 1.**
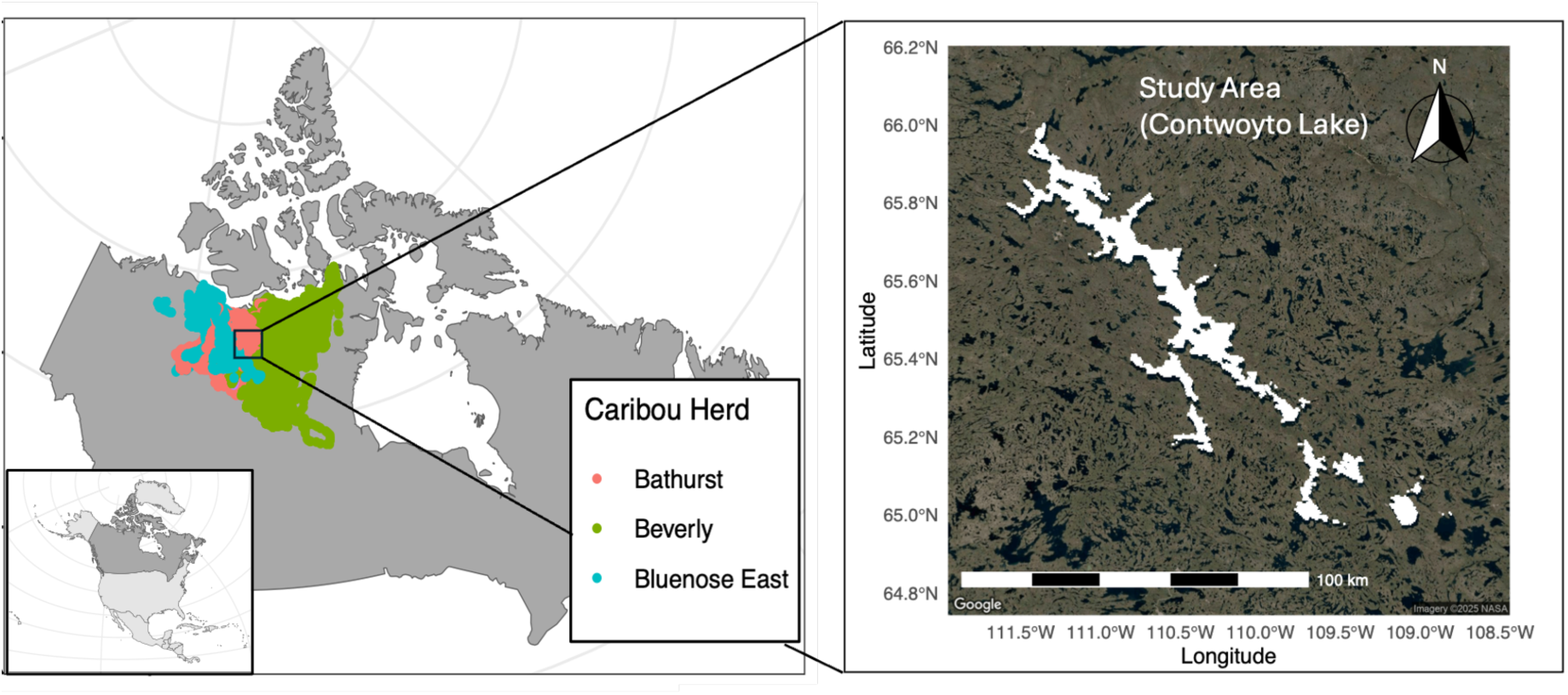
Location of the study area (Contwoyto Lake) in North America. The inset map in the bottom left corner of the left panel highlights Canada (in dark grey) within the broader context of North America. The left panel displays the historical migration ranges (2001–2021) of caribou herds who use areas around Contwoyto Lake: Bluenose East (blue), Bathurst (pink) and Beverly (green). These ranges are based on GPS tracking data from 406 individual barren-ground caribou (*Rangifer tarandus spp*.) The right panel highlights Contwoyto Lake, shown in the white area based on satellite imagery.

### 2.2 Animal Movement Data

We analyzed GPS tracking data from 406 adult barren-ground caribou whose movements occurred within 50 km of Contwoyto Lake between 2001 and 2021. The collars were deployed by the Government of the Northwest Territories’ Department of Environment and Climate Change (GNWT ECC) as part of ongoing population monitoring (Government of the Northwest Territories 2019). The dataset includes individual attributes such as sex, herd affiliation, unique ID, study site, timestamp, and geographic coordinates (longitude and latitude).

GPS fix rates varied both within and among individuals, ranging from 1-minute to 5-day intervals. Most data were collected at 8-hour intervals, with some 1-hour fixes triggered when animals entered geofenced zones around mining infrastructure. Occasional data gaps of up to one year were observed, likely due to signal loss or collar malfunction. Because our analysis focuses on the spatial interactions with lake ice conditions at fine temporal scales, we retained the original fix rates and did not filter or resample the data to preserve temporal heterogeneity and avoid introducing bias or loss of behavioral resolution.

### 2.3 Environmental Remote Sensing Data

We used two MODIS remote sensing products to characterize environmental conditions relevant to caribou movement and lake ice dynamics: (1) the MODIS Albedo Model (MCD43A3.061) and (2) the MODIS Land Cover Type product (MCD12Q1.061), both at 500 m resolution. The MCD43A3 Version 6.1 product provides daily estimates of shortwave broadband black-sky albedo (directional hemispherical reflectance) using 16-day composite data from both Terra and Aqua satellites from 2000 to 2023, offering high-quality surface reflectance for high-latitude regions (Schaaf et al. 2011). Because open water and frozen lake surfaces differ strongly in reflectance, albedo is particularly informative in this system as a remotely sensed indicator of changing lake surface conditions during freeze-thaw transitions. Due to polar night and low solar angle, however, albedo estimates between mid-October and late March are often unreliable. We therefore limited our analysis to the 92nd to the 280th day of the year (April 2 to October 7), when high-quality data are available (Appendix S1: Figure S1). Despite this limitation, the majority of the fall caribou migration near Contwoyto Lake is completed before October 7, ensuring that the methodology remains robust during the study period.

To identify lake boundaries, we extracted “pure water” pixels using the International Geosphere-Biosphere Programme - Data and Information System (IGBP-DIS) land cover classification (Loveland and Belward 1997; Belward et al. 1999; Sulla-Menashe and Friedl 2019) within the MCD12Q1.061 land cover dataset. This product provides annual global land cover types, with associated confidence assessments and quality control metrics from 2001 to 2021 (Sulla-Menashe and Friedl 2019). We adopted a multi-lake approach based on MODIS-derived pure water pixels rather than relying on administrative lake boundaries. This approach ensures that all remotely sensed variables used in our models, particularly albedo, are grounded in ecologically meaningful, observation-based definitions of lake surfaces. Methodological details and validation are provided in Appendix S1: Sections 3 and 4.

### 2.4 Processing Albedo Data

We use surface shortwave albedo as a proxy for lake ice conditions, leveraging its strong sensitivity to freeze-thaw transitions (Lucht et al. 2000). Albedo values are generally highest under frozen lake conditions and decline as melting progresses and open water emerges (Appendix S1: Section 5). To address data gaps due to cloud cover or sensor issues, we applied a gap-filling approach using a Kalman filter with spatial-temporal interpolation, following established climatological techniques (Welch and Bishop 1995; Jia et al. 2021; 2023; Appendix S1: Sections 6-8). These preprocessing ensured complete daily albedo time series for each pure water pixel of the lake across all study years.

To quantify the progression of ice melting and freezing, we introduced the “Albedo Percentile Rank” (APR) as a relative, pixel-specific measure of annual albedo position. For each pixel on each day of a given year, APR was calculated as the proportion of all albedo values observed at that pixel in that year that were less than or equal to the albedo value on that day. This index dynamically reflects a pixel’s position within its own annual albedo range, bounded by its year-specific maximum and minimum values, which we interpret as corresponding approximately to the most frozen and most thawed surface states observed for that pixel in that year. Calculated in this way, APR provides a localized, interannually standardized metric that captures continuous variation in ice conditions along the frozen-to-melted spectrum. For example, an APR of 90% indicates that the day’s albedo is higher than 90% of that pixel’s values in that year, corresponding to relatively early melt conditions. We used this spatially localized and year-specific measure because absolute albedo values can vary widely both among pixels within years and among years for a given pixel as a function of local environmental features, including lake depth, bottom topography, surface wetness, illumination geometry, and broader interannual variation in surface conditions (Perovich et al. 2002; Gardner and Sharp 2010; Fitzpatrick et al. 2014; Warren 2019; Leppäranta 2023). The use of APR does not imply that caribou respond to relative percentiles. Rather, APR provides a consistent, pixel-specific way to align each year’s ice conditions along a comparable frozen-to-melted progression, especially when absolute albedo values are influenced by spatial heterogeneity and interannual variability.

### 2.5 Processing Movement Data

To investigate how ice conditions affect caribou water-crossing behavior, we classified movement events near Contwoyto Lake into 3 categories: “Crossing Events”, “Circumnavigating Events”, and “Unknown Events”. These classifications were based on GPS locations and their spatial relationship to lake boundaries defined by MODIS-derived “pure water” pixels. “Crossing Events” were identified when at least one GPS fix from an individual’s track fell within the lake surface boundaries. These cases were treated as confirmed crossings, either over ice or open water, depending on season and ice conditions. “Circumnavigating Events” occurred when an individual’s track fell within the general vicinity of the lake and intersected with the virtual median line of the lake’s long axis, but no GPS points or interpolated segments crossed into the lake boundaries. We interpreted these as individuals went around the lake without crossing it.

“Unknown Events” were more ambiguous. These included cases where no GPS fix was recorded that occurred within the lake itself, but the line segments connecting consecutive GPS points did intersect the lake boundaries. Each classification was manually verified to reduce misclassification due to lake shape complexity or GPS error. Information regarding the number of events across different GPS fix intervals and the influence of temporal resolution on classification uncertainty is provided in Appendix S3: Section 1.

For each “Unknown Event”, we reconstructed two potential movement routes (Figure 2) across Contwoyto Lake: (1) a “Direct Path” defined as the shortest straight-line route between two consecutive GPS locations on opposite shores, and (2) a “Circumnavigate Path,” defined as the shortest overland route between the same points that avoids all open water. To further contextualize the decision, we extracted a “Reference Path,” consisting of the three on-land steps before and after the transit event, to estimate baseline movement behavior. All Circumnavigate Paths were generated using the *gdistance* R package (van Etten 2017), which applies a raster-based implementation of Dijkstra’s algorithm to approximate least-cost overland routes (Adriaensen et al. 2003; Etherington 2016). Because space is discretized, these paths represent close approximations rather than mathematically exact shortest paths. However, at the spatial resolution and extent of our analysis, this approximation is sufficient for characterizing relative circumnavigation costs (see Appendix S2). We derived three types of average speeds for each path (“Direct Speed”, “Circumnavigate Speed”, “Reference Speed”) due to varied GPS fix rates. Additional details on the classification of transit events, least cost path modeling, and path construction appear in Appendix S2, and full definitions of speed in Appendix S3: Section 2-3.

**Figure 2.**
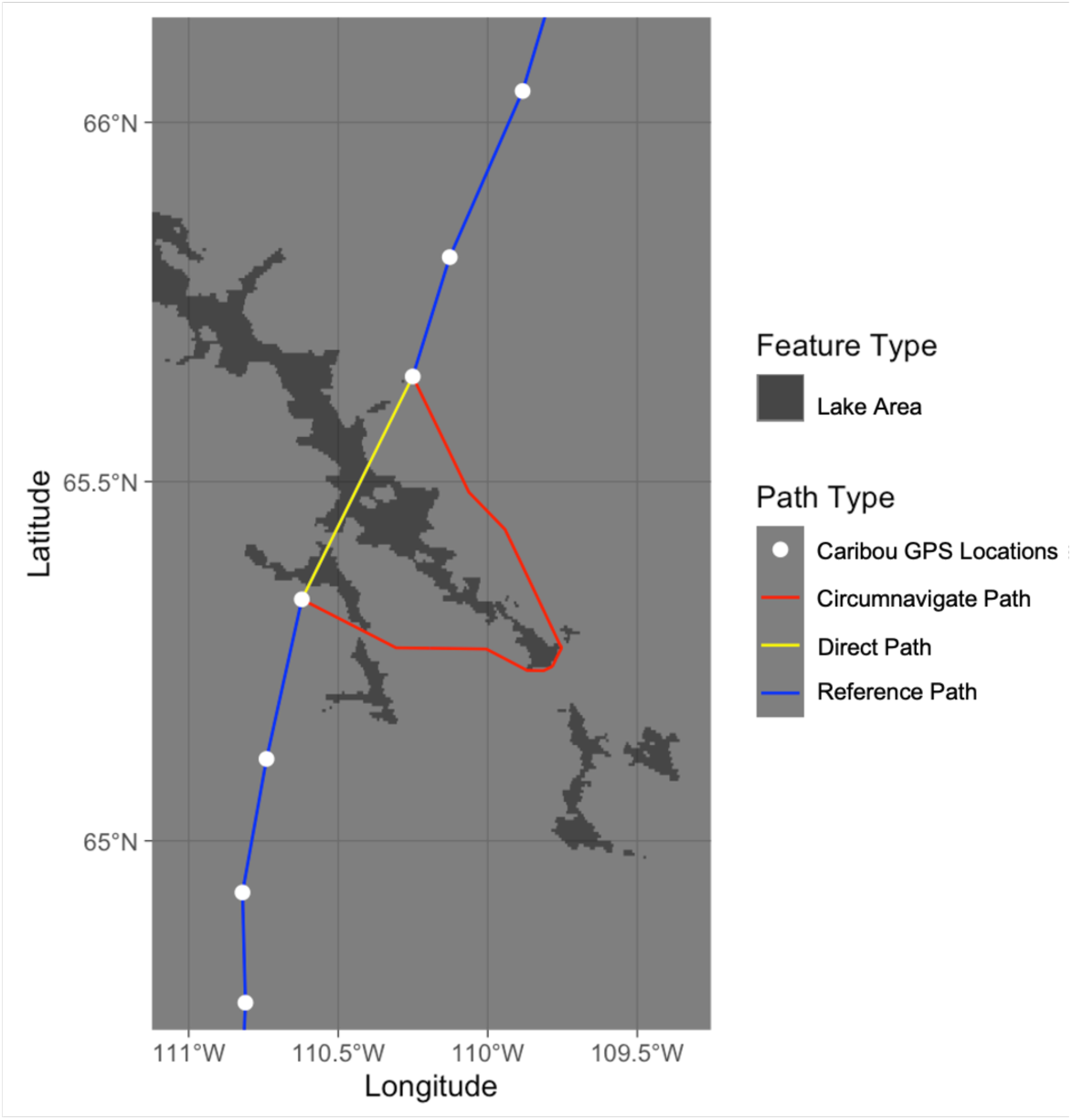
The direct path (yellow) and circumnavigate path (red) with the shortest distance between two consecutive starting and ending points (white) on the opposite side of the pure water areas of the Contwoyto Lake (dark grey) for one Unknown transit event (ID ‘bat106’ in 2012, around 2012-05-14 15:00:00) and the reference path (blue) shows 3 steps before and after the crossing.

To classify each “Unknown Event” as either a crossing or circumnavigating event, we trained two classification models, Random Forest (RF) and Binomial Logistic (BL) models, using “Crossing Events” and “Circumnavigating Events” as labeled training data (Venables and Ripley 2002; Breiman 2001). We retained both approaches because they differ in model assumptions and flexibility. The RF model provides a flexible nonparametric comparison that can accommodate nonlinearities and interactions, whereas the BL model provides a more interpretable parametric framework. Agreement between the two approaches therefore provides a useful check on the robustness of inferred classification patterns. These models incorporated ten predictor variables. Unless otherwise stated, all ice-related predictors described below use albedo percentile rank (APR), rather than absolute albedo values. Specifically, predictors spanned movement dynamics (Circumnavigate Speed / Reference Speed, Direct Speed / Reference Speed, and Circumnavigate Speed / Direct Speed), ice conditions at three spatial scales (APR at the Nearest Pixel, APR Along the Potential Crossing Path, and APR of the Entire Lake Area), lake features (Lake Width, Proportion of Time Spent in the Lake Area), and individual traits (Sex, Migration Season) (see Appendix S3: Section 4). Migration season was defined based on observed movement patterns, with “Spring Migration” occurring before July 1st, whereas “Fall Migration” occurred afterward. Because caribou movement characteristics differ markedly between spring and fall migration periods, we first conducted season-specific analyses by dividing the “Unknown Event” dataset into a “Spring Only” and a “Fall Only” subset. However, event types were unevenly distributed between these seasons, with “Crossing Events” dominating in spring and “Circumnavigating Events” dominating in fall, resulting in opposite forms of class imbalance in the seasonal datasets. To complement the season-specific analyses and mitigate these imbalances, we additionally constructed a “Combined” dataset that pooled events from both spring and fall migrations. This combined dataset provided a more balanced representation of event types and allowed us to evaluate whether predictors identified in seasonal analyses retained explanatory power when analyzed jointly.

For each dataset, 80% of the known events were randomly selected for model training, and the remaining 20% were reserved for testing. Model performance was evaluated using multiple metrics, including Accuracy, Sensitivity, F1 score, Cohen’s Kappa, and Area Under the Receiver Operating Characteristic Curve (AUC). For RF models, out-of-bag (OOB) error was recorded, whereas model selection for BL models was based on the delta Akaike Information Criterion (ΔAIC) (Appendix S3: Section 5). All predicted Unknown Events were labeled as “Crossing” or “Circumnavigating” based on a 50% predicted probability threshold.

To mitigate seasonal imbalance in event types (e.g., more crossing events in spring, more circumnavigating events in fall), we applied cost-sensitive weighting in the RF model (see details in Appendix S3: Section 6), and variable importance was measured by the Gini Importance Index. For the BL model, we applied Lasso regression (the *glmnet* R package) to penalize and exclude redundant predictors. Multicollinearity was assessed using Variance Inflation Factors (VIF), and redundant variables were removed when VIF > 5. After confirming the final predictors, we used binned residual plots to assess linearity and introduced quadratic terms only when U- or inverted U-shaped patterns were evident. Final selection was based on the best converged model with the lowest AIC using the *dredge* function (the *MuMIn* R package), retaining only predictors with P value < 0.05 as determined by the *glm* function (the *stats* R package). Model accuracy and classification were assessed using confusion matrices generated by the *confusionMatrix()* function from the *caret* R package and from the *randomForest* R package. ROC curves and AUC values were calculated using the *pROC* package to evaluate the models’ ability to discriminate between crossing and circumnavigating events.

In addition to the event-classification analyses above, we conducted a supplementary lake-scale analysis to place the spring crossing threshold identified from the behavioral analysis in a broader climatic context (Appendix S6). Specifically, after estimating the spring threshold from the classification analysis, we summarized lake-wide, pixel-level albedo values associated with the corresponding reference quantile and identified the first day in each year on which each pixel’s APR fell below that relative threshold. This supplementary analysis was designed to evaluate whether threshold-relevant ice conditions were characterized primarily by monotonic directional change or by strong interannual variability.

## 3. Results

Contwoyto Lake exhibited a clear seasonal cycle in surface conditions, as reflected in the APR pattern (e.g., in 2019; Figure 3C). APR values decreased from April (dark red), reached a minimum around September (dark blue), and then increased again by early October (light blue), reflecting the sequential process of ice melt followed by refreezing in the lake-wide average APR trend (Figure 3B). To further characterize spatial variation in ice phenology, we identified the first day each year that pixel-level albedo reached defined percentiles between its annual maximum and minimum. This provided spatially explicit estimates of ice melt and freeze timing during the spring and fall migration seasons (see Appendix S4: Figure S1-S3)

**Figure 3.**
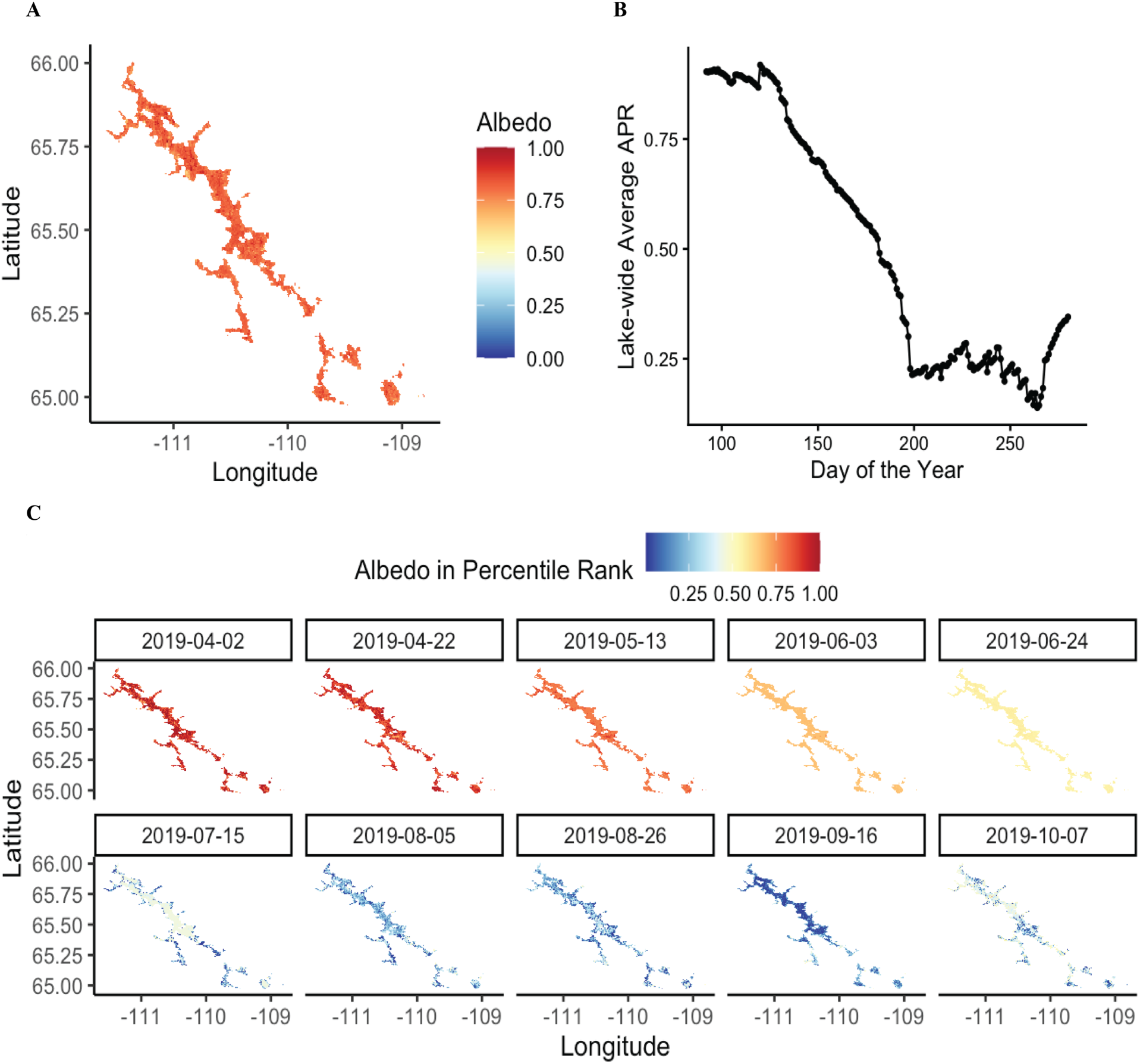
Processing of the albedo data. (A) Albedo of Contwoyto Lake on the 92nd day (April 2nd) in 2019 after preprocessing. Data gaps were filled using temporal-spatial interpolation with climatology and a Kalman filter, and pure water bodies were extracted using land cover data. (B) Lake-wide average Albedo Percentile Rank (APR) from April 2nd (92nd day) to October 7th (280th day) in 2019. (C) Spatial distribution of Albedo Percentile Rank (APR) for pure water body pixels from April 2nd (dark blue) to October 7th (dark red) in 2019.

In total, we identified 367 transit events from 130 individuals, all occurring between 2005 and 2021. Of these, 145 occurred during spring migration and 222 during fall migration. In spring migration, 72 cases were classified as known events (59 crossing, 13 circumnavigating), and 73 cases were classified as unknown. In fall migration, 97 cases were known events (3 crossing, 94 circumnavigating), and 125 were unknown (see Table 1)

**Table 1.**
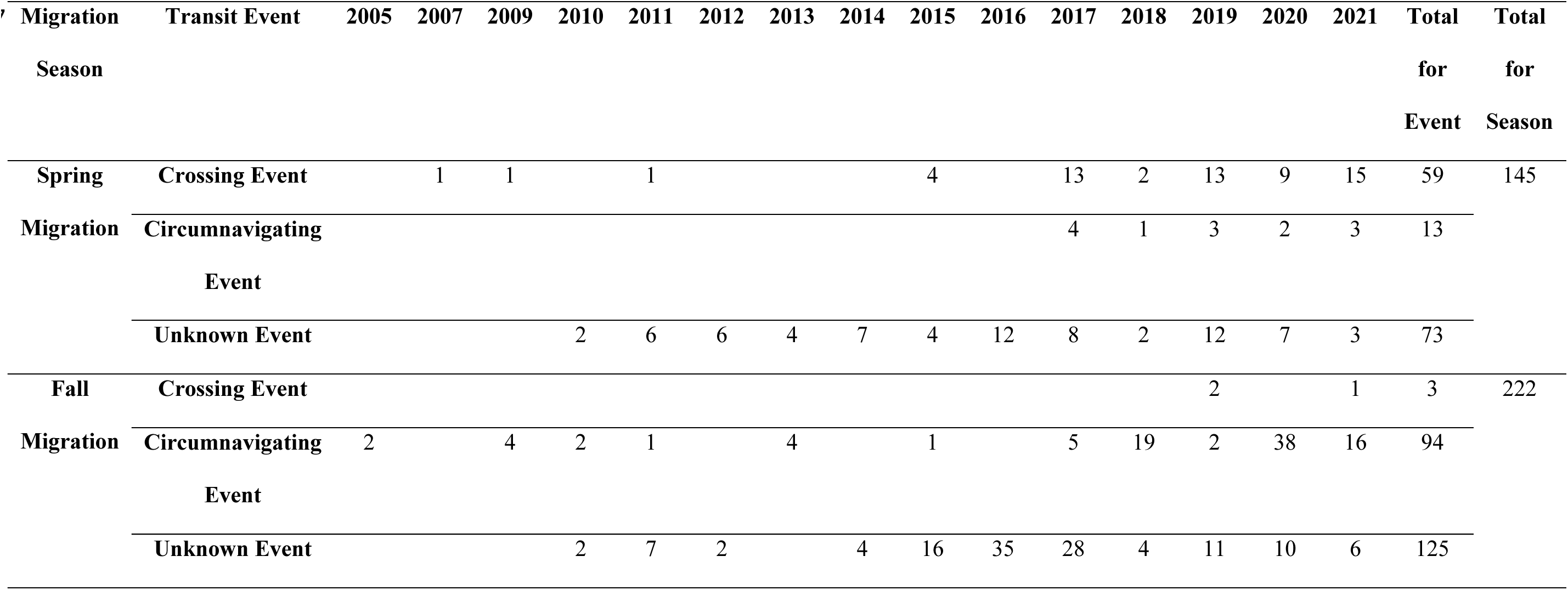
Distribution of caribou transit events across Contwoyto Lake by event type and migration season (2005–2021), illustrating both seasonal patterns and interannual variability in water-crossing behavior, potentially shaped by behavioral plasticity or limitations in detection.

| 347 | Migration | Transit Event | 2005 | 2007 | 2009 | 2010 | 2011 | 2012 | 2013 | 2014 | 2015 | 2016 | 2017 | 2018 | 2019 | 2020 | 2021 | Total | Total |
| --- | --- | --- | --- | --- | --- | --- | --- | --- | --- | --- | --- | --- | --- | --- | --- | --- | --- | --- | --- |
|  | Season |  |  |  |  |  |  |  |  |  |  |  |  |  |  |  |  | for | for |
|  |  |  |  |  |  |  |  |  |  |  |  |  |  |  |  |  |  | Event | Season |
|  | Spring | Crossing Event |  | 1 | 1 |  | 1 |  |  |  | 4 |  | 13 | 2 | 13 | 9 | 15 | 59 | 145 |
|  | Migration | Circumnavigating |  |  |  |  |  |  |  |  |  |  | 4 | 1 | 3 | 2 | 3 | 13 |  |
|  |  | Event |  |  |  |  |  |  |  |  |  |  |  |  |  |  |  |  |  |
|  |  | Unknown Event |  |  |  | 2 | 6 | 6 | 4 | 7 | 4 | 12 | 8 | 2 | 12 | 7 | 3 | 73 |  |
|  | Fall | Crossing Event |  |  |  |  |  |  |  |  |  |  |  |  | 2 |  | 1 | 3 | 222 |
|  | Migration | Circumnavigating | 2 |  | 4 | 2 | 1 |  | 4 |  | 1 |  | 5 | 19 | 2 | 38 | 16 | 94 |  |
|  |  | Event |  |  |  |  |  |  |  |  |  |  |  |  |  |  |  |  |  |
|  |  | Unknown Event |  |  |  | 2 | 7 | 2 |  | 4 | 16 | 35 | 28 | 4 | 11 | 10 | 6 | 125 |  |

### Crossing Events

A total of 109 GPS relocations from 47 individuals occurred within the boundaries of Contwoyto Lake. These relocations were unevenly distributed, with more events observed during spring migration. Summary statistics, including crossing dates, average APR on the day of crossing, and landscape type where the relocations were located (i.e., lake water or lake island), are presented in the Appendix S4: Figure S4. During spring migration, direct crossing paths showed a consistent southwest-to-northeast orientation (Figure 4A, B). In contrast, only a few crossing events were recorded in fall migration (Figure 4E, F). These fall events featured narrower and more variable trajectories, reflecting a more spatially constrained and possibly more cautious movement pattern.

**Figure 4.**
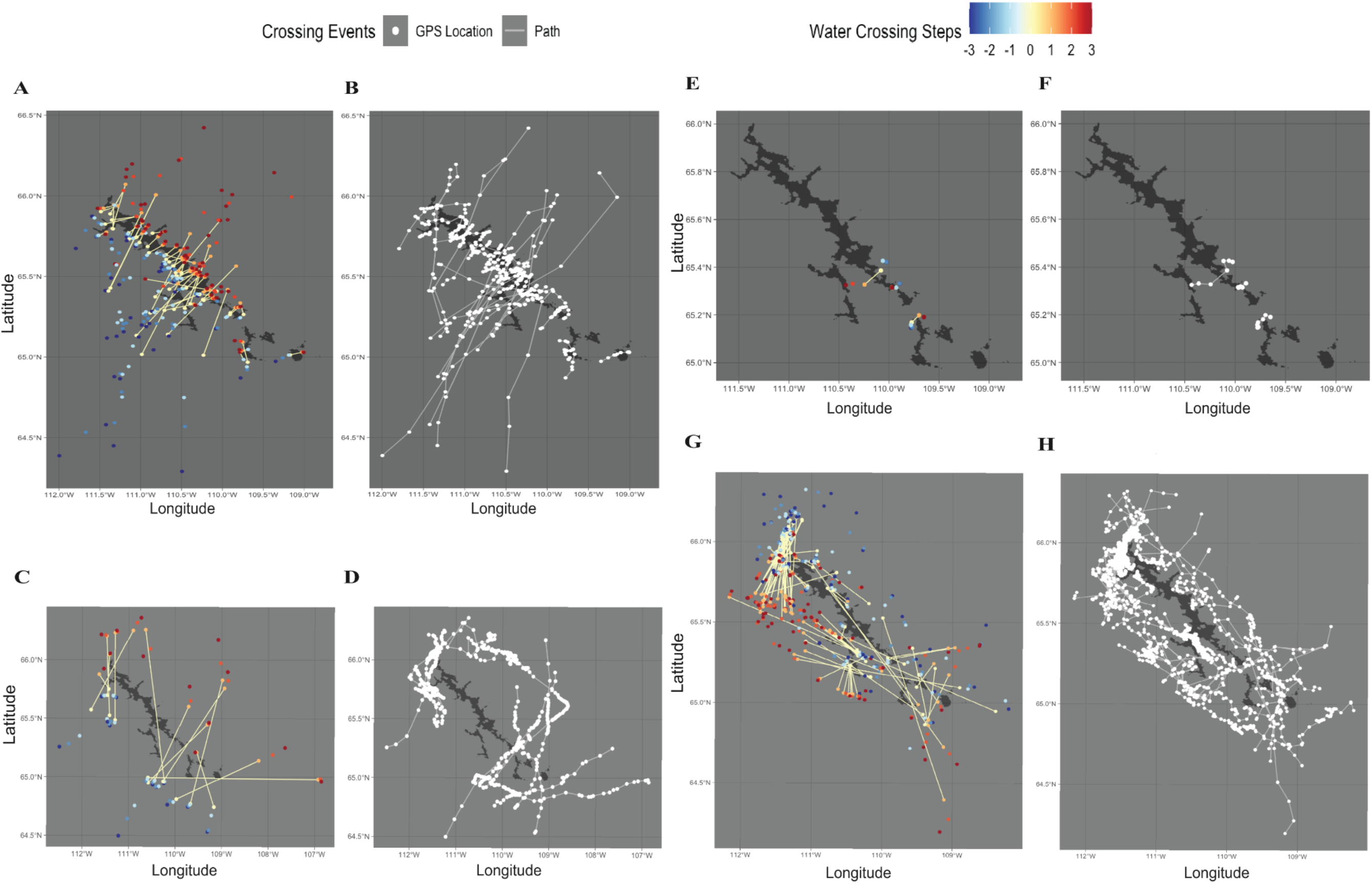
Spatial Distribution of Caribou trajectories for known events (2001–2021). Event Types: Crossing (A, B, E, F), Circumnavigating (C, D, G, H), during spring (A—D) and fall (E—H) migrations. Spring Migration: (A, B) 59 crossing events; (C, D) 13 circumnavigating events. Fall Migration: (E, F) 3 crossing events; (G, H) 94 circumnavigating. (A, C, E, G) shows direct paths of transit events with the starting (yellow) and ending (orange) points, with steps before/after transit shown from -3 (dark blue) to +3 (dark red). (B, D, F, H) shows trajectories of the transit events (white).

### Circumnavigating Events

Using a 20 km buffer around Contwoyto Lake, we identified 13 circumnavigating events from 12 individuals during spring migration. These events lasted between 2 and 40 days, with potential cross-lake distances ranging from 0.33 to 29.3 km. In fall, 94 circumnavigating events were recorded from 57 individuals with durations between 0.16 and 65 days and the potential cross-lake distances from 0.93 to 47.9 km. In spring migration, circumnavigating routes were distributed throughout the lake: 5 individuals bypassed the northern portion, 4 the central region, and 4 the southern part (Figure 5C, D). Most of these paths were west-to-east, similar to the pattern observed in crossing events. In contrast, fall migration exhibited more frequent and spatially diverse circumnavigations (Figure 5G, H), with less directional consistency compared to spring (Figure 5C).

**Figure 5.**
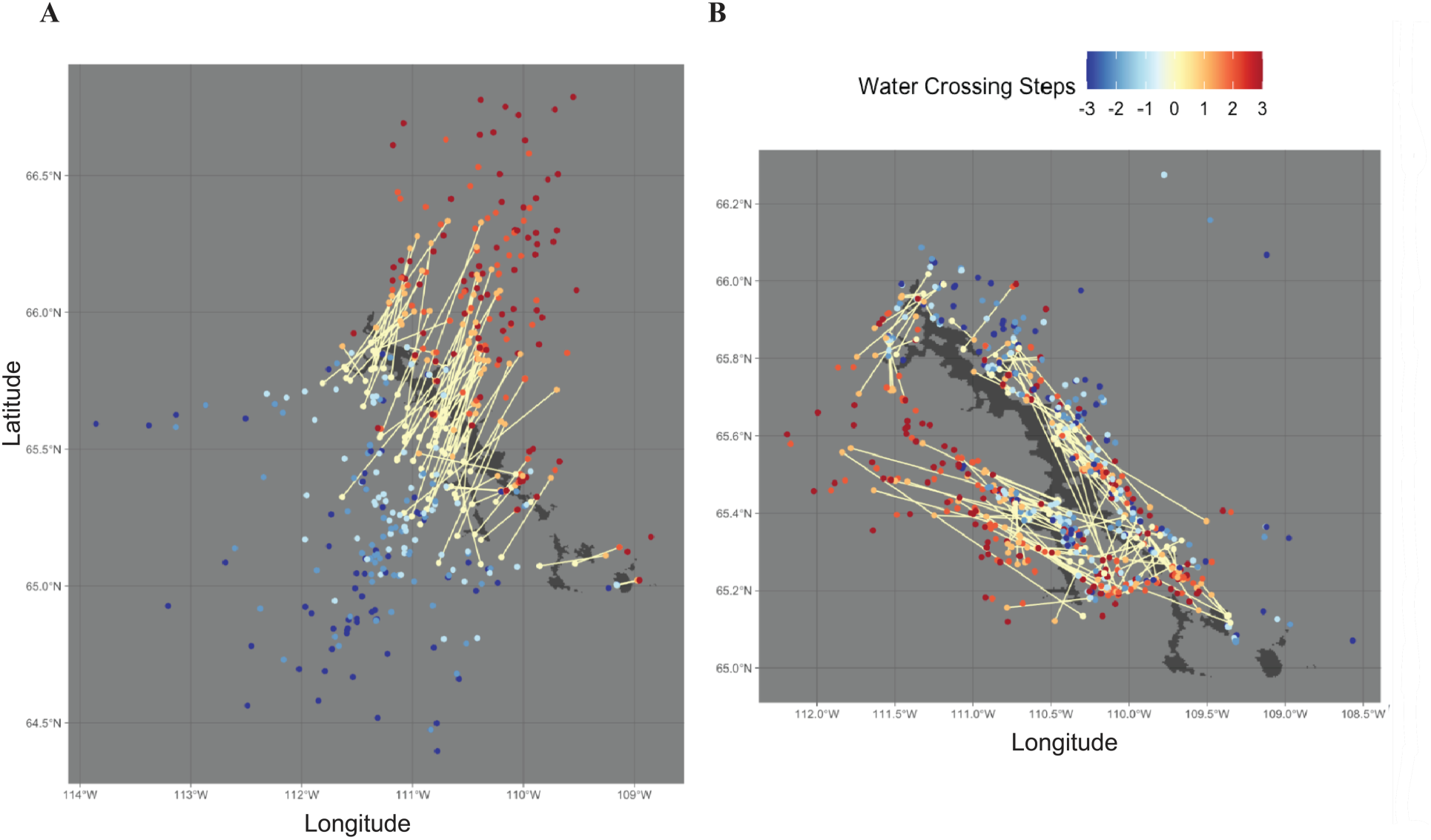
Spatial Distribution of Caribou trajectories for 198 unknown events (2001–2021). (A) Spring migration (73 unknown events); (B) Fall migration (125 unknown events). Both show direct paths of transit events with the starting (yellow) and ending (orange) points, with steps before/after transit shown from -3 (dark blue) to +3 (dark red).

### Unknown Events

A total of 73 unknown events were identified in spring migration and 125 during fall migration. In spring, nearly all direct paths followed a southwest-to-northeast direction (Figure 5A). In fall, however, direct paths showed more diverse orientations and were often aligned with or parallel to the lakeshore (Figure 5B).

Comparing the most important variables across RF and BL models, “APR Along the Potential Crossing Path” consistently emerged as a key predictor in the “Spring Only” dataset, underscoring the central role of ice conditions in shaping water-crossing behavior when lakes are not fully thawed in spring migration, before July 1st. In contrast, during fall migration, when the lake ice is almost ice-free, albedo loses its predictive value. Instead, movement-related metrics, such as speed ratios and the proportion of time spent in lake areas, became dominant. Notably, in the “Combined” Dataset, albedo regained importance alongside movement dynamics, suggesting that ice conditions and behavioral flexibility jointly influence caribou crossing decisions. Detailed analyses supporting these findings are discussed below.

In the “Spring Only” dataset, both the RF and BL models demonstrated high classification accuracy. The RF model achieved perfect performance on the test data (Accuracy = 1.00; 95% CI: 0.782–1.00; Kappa = 1.00; F1 = 1.00), with a low OOB error. According to the Gini Importance Index (Figure 6A), the top three predictors were: (1) APR Along the Potential Crossing Path (medium scale), (2) The Ratio of Circumnavigate Speeds Over Direct Speeds (log), and (3) The Ratio of Circumnavigate Speeds Over Reference Speeds (log). The best-performing BL model retained “APR Along the Potential Crossing Path” as the only significant predictor (β = 43.24, SE = 21.29, P = 0.0422). Predicted crossing probability increased with albedo, with a 50% crossing probability corresponding to an albedo percentile rank of 0.56 (Figure 6B).

**Figure 6.**
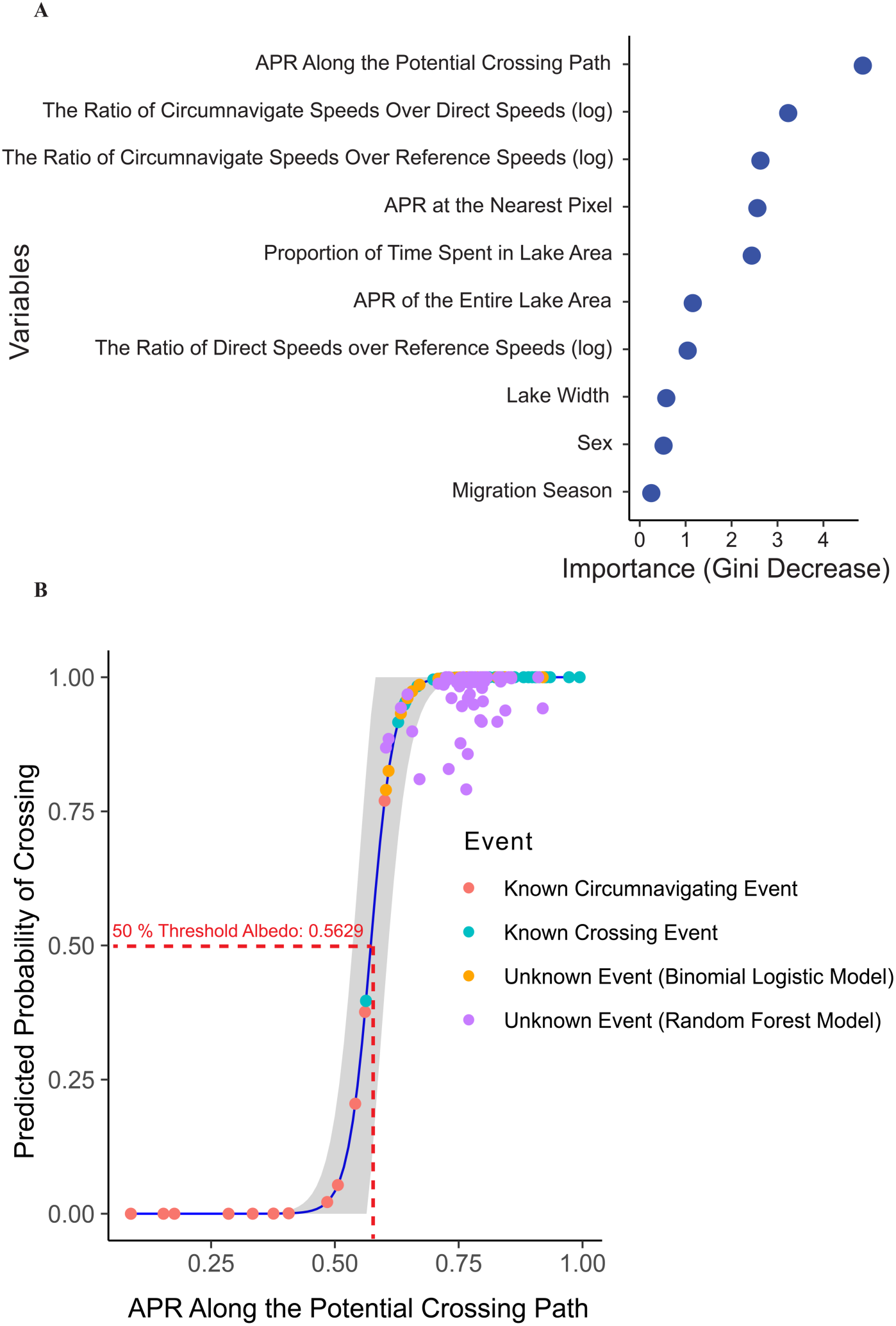
Model fitting results for the Random Forest (RF) and Binomial Logistic (BL) models in the “Spring Only” dataset. (A) Gini Importance Index (also called Mean Decrease Gini Score) from the RF model classifying spring transit events (crossing vs. circumnavigating). (B) BL model showing crossing probability increasing with higher APR along potential crossing paths (blue line with 95% CI in grey), trained on Crossing (green points) and Circumnavigating Events (red points). Both models (RF results in purple points, BL results in yellow points) predicted crossing probabilities above 50% for Unknown Events, indicating strong agreement in classifying caribou water crossing behaviors during spring migration.

In the “Fall Only” dataset, model performance was affected by extreme class imbalance (3 crossings vs. 94 circumnavigating events). Although the RF model achieved high overall accuracy (95%, 95% CI: 0.75–0.99), it failed to correctly identify any crossing events (Sensitivity = 0, F1 = NA, Kappa = 0), highlighting the limitations of the model under skewed data (Table 2). The BL model performed relatively better, correctly identifying some crossing events (Sensitivity = 66.67%). In this dataset, the most influential predictor in the BL model was “The Ratio of the Circumnavigate Speeds Over Reference Speeds” (β = 2.6239, SE = 0.9725, P = 0.00694), with predicted crossing probability exceeding 50% when this ratio surpassed 2.698 (Appendix S5: Figure S1B). No albedo-related variables ranked among the top predictors in either model.

**Table 2.**
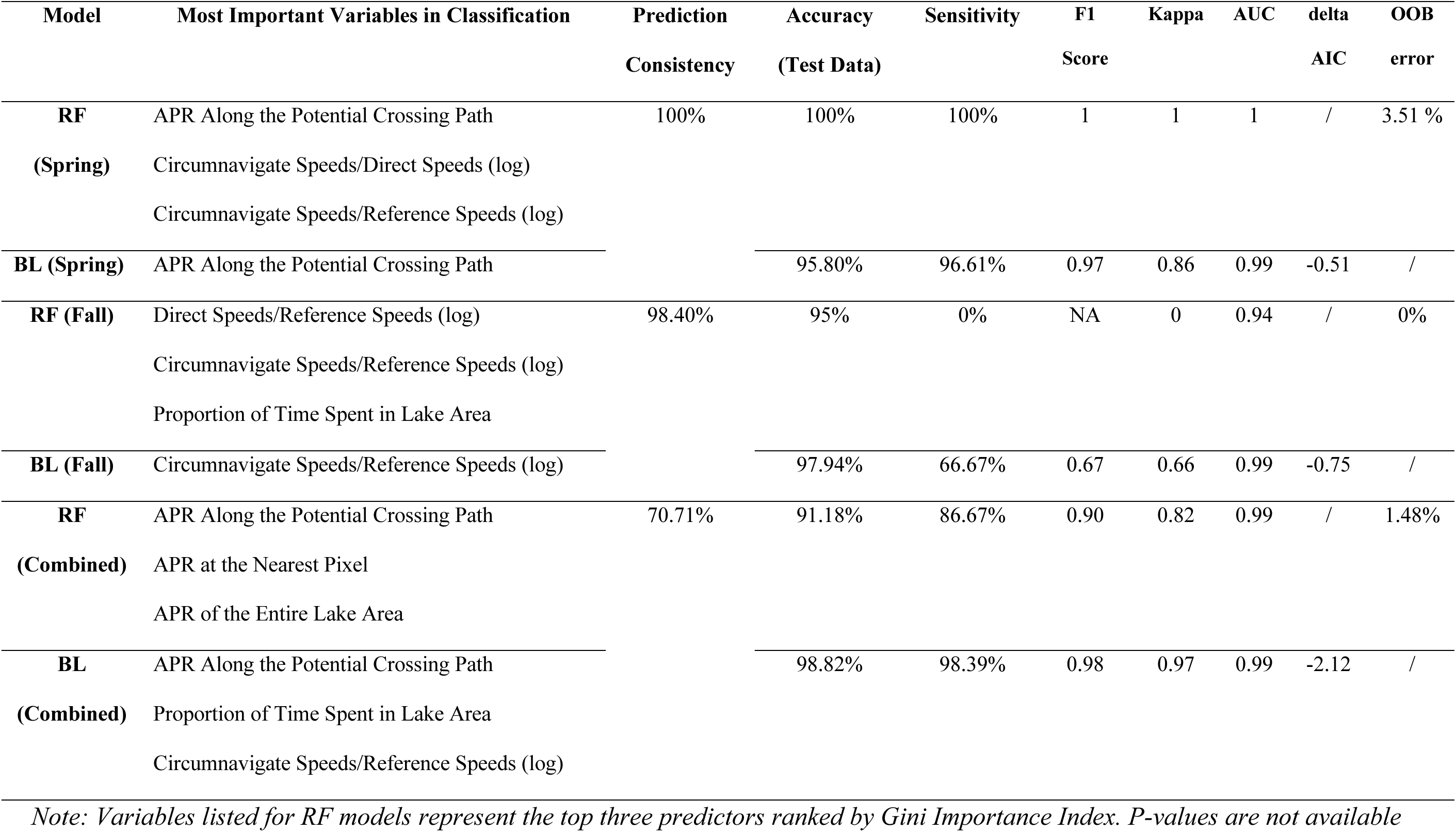
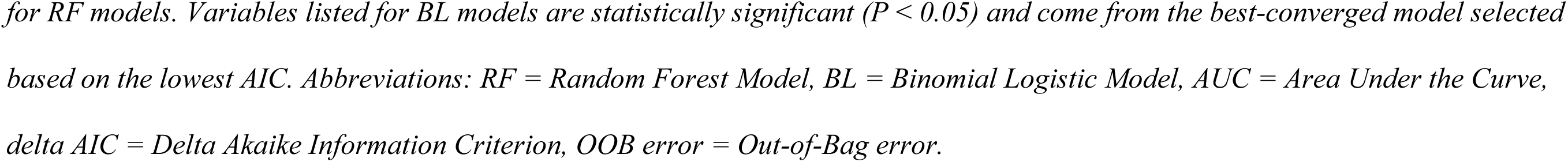
Comparison of model performance and key predictors for classifying caribou lake crossing behavior in “Spring Only”, “Fall Only”, and “Combined” datasets.

| Model | Most Important Variables in Classification | Prediction<br>Consistency | Accuracy<br>(Test Data) | Sensitivity | F1<br>Score | Kappa | AUC | delta<br>AIC | OOB<br>error |
| --- | --- | --- | --- | --- | --- | --- | --- | --- | --- |
| **RF**<br>**(Spring)** | APR Along the Potential Crossing Path | 100% | 100% | 100% | 1 | 1 | 1 | / | 3.51 % |
|  | Circumnavigate Speeds/Direct Speeds (log) |  |  |  |  |  |  |  |  |
|  | Circumnavigate Speeds/Reference Speeds (log) |  |  |  |  |  |  |  |  |
| **BL (Spring)** | APR Along the Potential Crossing Path |  | 95.80% | 96.61% | 0.97 | 0.86 | 0.99 | -0.51 | / |
| **RF (Fall)** | Direct Speeds/Reference Speeds (log) | 98.40% | 95% | 0% | NA | 0 | 0.94 | / | 0% |
|  | Circumnavigate Speeds/Reference Speeds (log) |  |  |  |  |  |  |  |  |
|  | Proportion of Time Spent in Lake Area |  |  |  |  |  |  |  |  |
| **BL (Fall)** | Circumnavigate Speeds/Reference Speeds (log) |  | 97.94% | 66.67% | 0.67 | 0.66 | 0.99 | -0.75 | / |
| **RF**<br>**(Combined)** | APR Along the Potential Crossing Path | 70.71% | 91.18% | 86.67% | 0.90 | 0.82 | 0.99 | / | 1.48% |
|  | APR at the Nearest Pixel |  |  |  |  |  |  |  |  |
|  | APR of the Entire Lake Area |  |  |  |  |  |  |  |  |
| **BL**<br>**(Combined)** | APR Along the Potential Crossing Path |  | 98.82% | 98.39% | 0.98 | 0.97 | 0.99 | -2.12 | / |
|  | Proportion of Time Spent in Lake Area |  |  |  |  |  |  |  |  |
|  | Circumnavigate Speeds/Reference Speeds (log) |  |  |  |  |  |  |  |  |

*Note: Variables listed for RF models represent the top three predictors ranked by Gini Importance Index. P-values are not available*
*for RF models. Variables listed for BL models are statistically significant ( $P < 0.05$ ) and come from the best-converged model selected* *based on the lowest AIC. Abbreviations: RF = Random Forest Model, BL = Binomial Logistic Model, AUC = Area Under the Curve,* *delta AIC = Delta Akaike Information Criterion, OOB error = Out-of-Bag error.*

The “Combined” dataset, which integrates both spring and fall events, yielded strong model performance for both approaches. The RF model achieved 91.2% accuracy on the test data (95% CI: 0.76–0.98), outperforming the No Information Rate (P = 1.9e–06). The three most influential predictors were all albedo-related: “APR Along the Potential Crossing Path”, “APR at the Nearest Pixel”, and “APR of the Entire Lake Area” (Appendix S5: Figure S1C). The best BL model achieved 98.8% accuracy and identified three significant predictors: “APR Along the Potential Crossing Path” (β = 13.78, SE = 3.98, P < 0.001), “Proportion of Time Spent in Lake Area” (β = 19.63, SE = 9.95, P = 0.049) and “The Ratio of Circumnavigate Speeds Over Reference Speeds (log)”(β = 1.83, SE = 0.83, P = 0.027). Three crossing events in the “Combined” dataset occurred at low values of “APR Along the Potential Crossing Path” (with percentile rank < 0.46), all of which occurred during fall migration (Appendix S5: Figure S1D). Summary statistics for agreement between RF and BL classifications of unknown events are provided in Table 2 and Appendix S5: Sections 1-3.

To place the behaviorally derived threshold identified in the spring-only BL model in broader lake-scale climatic context, we quantified interannual variability in lake-wide, pixel-level 56th-percentile albedo values and identified the timing at which each pixel’s albedo percentile rank (APR) first dropped below its annual 56th percentile during the spring melt-down period each year from 2005 to 2021 (Appendix S6: Figure S1, Table S1). Neither the annual 56th percentile albedo value nor the timing of this threshold exhibited a strong monotonic temporal trend. Linear trend analyses detected statistically significant but ecologically negligible shifts, corresponding to changes of less than half a day over the study period, with “Year” explaining less than 1% of the total variance in both albedo values and threshold-crossing timing (R² < 0.01). Instead, ice conditions relative to the behavioral threshold were dominated by pronounced interannual variability. When “Year” was treated as a categorical factor, models explained 65% of the variance in albedo values and 12% of the variance in threshold-crossing timing. Certain years reached threshold conditions substantially earlier than others (e.g., 2018), whereas late-threshold years (e.g., 2020) exhibited delayed transitions. Together, these results suggest that, over the 2005–2021 study period, variation in lake ice conditions surrounding the behaviorally derived albedo threshold was expressed primarily through strong interannual volatility, with little evidence for a monotonic long-term trend (see Appendix S6 for details).

## 4. Discussion

By aggregating 20 years of GPS tracking data with daily MODIS albedo observation, we developed a novel, scalable framework linking seasonal lake ice conditions to caribou water-crossing behavior. Our results reveal a clear seasonal shift in behavioral drivers: in spring, surface albedo, as a proxy for ice condition, best predicted crossing behavior, while in fall, movement-based metrics became more influential, indicating that open water poses a greater barrier. These findings underscore how dynamic surface conditions act as seasonal behavioral filters that shape migratory strategies, offering mechanistic insights into species’ responses to climate-driven disruptions in landscape connectivity.

### 4.1 Using Albedo as an Indicator of Ice Condition

We developed a novel approach to quantify lake ice dynamics using MODIS-derived albedo, offering a rare combination of spatial resolution (500m), daily temporal frequency, and two decades of historical coverage. The method captures pixel-level freezing and melting patterns across large Arctic lakes and aligns seamlessly with long-term caribou GPS data, enabling inference about movement responses across both space and time. By using surface albedo as a proxy for ice conditions, our method overcomes the limitations of sparse in-situ monitoring (Magnuson et al. 2000; Eklund 1999) and provides essential environmental context for interpreting individual water-crossing decisions.

Compared to previous remote sensing approaches, our method strikes a critical balance between spatial, temporal, and historical dimensions. For example, while Leblond et al. (2016) used 8-day average NDSI, our daily product allows finer temporal alignment with animal movement. Synthetic Aperture Radar (SAR) based methods (Du et al. 2015; Stonevicius et al. 2022) provide greater spatial detail but suffer from coarse revisit frequency (5-12 days). Giroux-Bougard (2023) applied the OPEN-ICE algorithm to 30-m fused Sentinel-2 and Landsat7/8 imagery, but this approach is only available post-2013. Other lake ice products, such as the European Space Agency’s Lakes Climate Change Initiative (Crétaux et al. 2022) and Copernicus Global Land Service (CGLS) Lake Ice Extent (LIE) product (2022) offer daily coverage but at either coarser resolution (1000m) or over shorter time spans. Passive microwave sensors (Matias et al. 2024) offer broad coverage but at much coarse resolution (3 km), and lake-point methods (Šmejkalová et al. 2016) cannot resolve spatial heterogeneity within large lakes, an essential feature for linking conditions to animal decisions at migration-relevant scales.

In addition to providing improved data alignment, our method contributes to the standardization of ice condition metrics, an ongoing challenge in lake ice phenology research. Rather than relying on inconsistent definitions of “freeze” or “break-up” dates, we introduce a continuous, percentile-based metric derived from each pixel’s annual albedo range. This allows for a more nuanced characterization of ice transitions and accommodates spatial variability caused by lake depth, bathymetry, and microclimatic factors. Importantly, during spring melt, surface albedo can decline rapidly due to wet snow, slush, or meltwater films (e.g., melt ponds; Perovich et al. 2002; Warren 2019) while ice remains continuous and load-bearing. Under these conditions, absolute albedo becomes a poor indicator of functional permeability. By employing a pixel-specific, within-year normalization, the APR metric helps align ice conditions along a comparable frozen-to-melted progression, reducing interannual variability driven by surface moisture, illumination geometry, and sensor-related effects. Unlike thresholding approaches that classify binary ice-water states (Pavelsky and Smith 2004; Cooley et al. 2016), our method retains pixel-level details across the lake surface, enhancing alignment with animal movement and supporting behavioral modeling at intermediate spatial scales. Moreover, by tracking percentile shifts across years, this framework facilitates long-term monitoring of melt/freeze directionality and its implications for behavioral plasticity under climate change.

Despite these advantages, our method is constrained by Arctic-specific limitations inherent to optical remote sensing. MODIS albedo data are only reliable from April to early October (days 92–280), due to low solar angle and polar night conditions, limiting our ability to analyze late fall freeze-up. Additionally, while surface albedo is an effective proxy for ice conditions, it does not directly capture ice thickness or load-bearing capacity, which may ultimately determine crossing feasibility. The 500-m spatial resolution, though adequate for general spatial patterns, may overlook small-scale heterogeneity near lake edges or in narrow inlets that influence individual movement decisions. These constraints necessitated the exclusion of certain late-season events and limited our analysis of fall migration. Future improvements may involve integrating optical and active microwave (e.g., SAR) remote sensing, combined with machine learning approaches (Murfitt and Duguay 2021), to enhance spatial and temporal resolution across broader timeframes.

### 4.2 Water Crossing Behavior

Our results reveal how caribou adjust migratory behavior in response to lake ice conditions, with particularly strong signals in spring. Surface albedo, used as a proxy for ice condition, was the most influential predictor of crossing decisions. We identified a clear behavioral threshold: when APR along the crossing path exceeded the 56th percentile, caribou were more likely to cross than circumnavigate. Crossing probability rose to 70% at the 60th percentile and reached ∼90% at the 63rd percentile, while it dropped to 10% at the 51st percentile. This quantifiable, remotely observable threshold provides a concrete metric for understanding when rapid changes in landscape permeability alter migratory route choice.

In contrast, fall migration patterns were shaped primarily by the presence of open water. In the absence of lake ice, caribou relied on movement-based metrics, particularly the log-transformed ratio of circumnavigate speeds over reference speeds. This shift in behavioral drivers corresponded to a pronounced seasonal transition: crossing events dropped from 82% of spring observations to just 3% in fall, while circumnavigating events increased from 18% to 97%. These patterns underscore the role of lake ice as a seasonal enabler of movement and the extent to which open water poses a behavioral barrier to caribou (Leblond et al. 2016; Leclerc et al. 2021).

Notably, we identified three fall migration events (“bat244”, “bat292”, and “bat205”) likely involving caribou swimming across open water. These events occurred at narrow lake segments, likely traditional crossing corridors (Williams and Gunn 1982; Gordon 1977; 1996; 2003; 2005), and were characterized by low albedo values (<46th percentile) and slower speeds (∼ 0.39 m/s) compared to their spring over-ice crossings (∼ 0.50 m/s). The average swimming distances (∼ 1794 m) were consistent with prior field observations, such as a documented 1600 m swim by a caribou calf in saltwater (Miller 1995). Caribou are known strong swimmers, often motivated by food opportunities, predator avoidance, or insect harassment (Webber et al. 2021; Jeffery et al. 2007; Jordan et al. 2010). Even young calves follow their mothers across open water without hesitation (Skoog 1968). Reported swim speeds can reach 2.68 m/s during escape (Banfield 1961), suggesting that our observed values are more consistent with slow, non-escape crossings through open water than with high-speed escape responses.

Despite strong behavioral patterns, some uncertainty remains in classifying water crossing events. Our approach relied primarily on spatial analyses of GPS relocations relative to lake boundaries, complemented by visual inspection of movement trajectories. Without direct field validation, brief exploratory movements, such as shoreline hesitation, backtracking, or aborted crossings, may be misclassified. Commercial high-resolution satellite imagery offers a promising avenue for post hoc validation, potentially revealing movement traces, disturbance patterns, or even individual animals around suspected crossing sites (Fretwell et al. 2012; Stapleton et al. 2014; Duporge et al. 2020; Wu et al. 2023).

Additional uncertainty arises from variability in GPS fix rates. In some years, tracking occurred at coarse temporal resolutions (e.g., every 24 or even 48 hours), and fixes were occasionally missing during transit phases, particularly in late spring when ice conditions fluctuate rapidly. A supplementary analysis of “Unknown” events across GPS fix interval categories (Appendix S3: Section 1) indicates that coarser temporal resolution is associated with a higher proportion of unclassified events, suggesting that classification uncertainty is partly driven by sampling interval rather than behavioral ambiguity per se. GPS fix schedules in long-term tracking studies necessarily reflect trade-offs among battery capacity, deployment duration, and collar weight (Hebblewhite and Haydon 2010; Kays et al. 2015). These gaps may obscure short crossings or bias the detection of crossing versus circumnavigation behavior. Nonetheless, the strength and consistency of seasonal patterns in our dataset suggest that these limitations do not obscure the broader behavioral signal.

Our findings also underscore the broader ecological importance of ice as a facilitator of migratory connectivity. In other Arctic regions, sea ice similarly serves as a critical bridge for inter-island (Peary Caribou) and island-mainland (Dolphin-Union) migration (Miller et al. 2005; Jenkins et al. 2016). These caribou show high site fidelity to specific sea-ice corridors and often delay movement until stable surfaces form (Poole et al. 2010); early crossing can be fatal (Dumond et al. 2013). As climate change reduces the duration and reliability of ice cover, caribou may face higher energetic costs from longer detours (in our parlance, circumnavigating events) and increased mortality risks, particularly for vulnerable calves (Miller 1995; Miller et al. 2005; Dumond et al. 2013). This erosion of seasonal bridges may constrain migratory flexibility and reduce the availability of efficient seasonal routes across Arctic landscapes.

Our results also raise new questions about the spatial and cognitive scales at which caribou perceive and respond to ice conditions. The superior predictive performance of crossing-path albedo compared to local (nearest-pixel) or lake-wide (entire lake) measures suggests that caribou respond to environmental conditions at intermediate spatial scales, potentially reflecting perceptual constraints in dynamic landscapes (Zollner and Lima 1997; Fagan et al. 2017). Prior experience may also contribute to these responses, potentially through experienced individuals guiding herds along historically safer routes (Williams and Gunn 1982; Jesmer et al. 2018; Merkle et al. 2019). Future research that incorporates individual age, sex, and memory-based behavior will improve our understanding of how environmental cues and past experience jointly shape migratory decision-making in complex and variable Arctic landscapes (Fagan et al. 2013; Bracis and Mueller 2017; Gurarie et al. 2021).

### 4.3 Migration Under Global Climate Change

Arctic warming exemplifies a broader class of climate-driven disruptions to seasonal connectivity (Post et al. 2009; Serreze and Barry 2011; Zeigler and Fagan 2014), with lake ice phenology offering a particularly acute case in northern systems. Earlier break-up and later freeze-up (Adrian et al. 2009; Dibike et al. 2012) have significantly reduced the seasonal permeability of safe crossing surfaces. These shifts directly affect migratory connectivity in water-rich northern landscapes (Downing et al. 2006). While prior studies have emphasized the timing of migration and arrival at calving grounds (Gurarie et al. 2019; Matias et al. 2024), our results provide finer-grained behavioral insight into how decisions are shaped by climate-sensitive barriers. Specifically, we identify quantifiable albedo thresholds that predict spring crossings, offering a behavioral indicator of environmentally suitable crossing conditions. As such, the proportion of crossing versus circumnavigating events may serve as a sensitive behavioral indicator of changing movement constraints in response to small-scale habitat change.

Interpreting these indicators requires accounting for the spatial complexity of the frozen landscape. Importantly, the timing of lake-wide mean ice melt should not be interpreted as the sole constraint on migratory decision-making. Contwoyto Lake is a long (>110 km), narrow (<10 km) northwest–southeast oriented system that spans a significant latitudinal range. Consequently, ice melt progresses heterogeneously along this gradient. Our pixel-level analysis (Appendix S6: Figure S1B) reveals that while the lake-wide mean break-up often occurs later (DOY 155–180), localized ice degradation in specific corridors begins significantly earlier, with the lower tail of the distribution reaching the 56th percentile threshold as early as DOY 141 (e.g., May 20). Notably, this onset of localized deterioration is temporally closer to the end of the spring migration window described by Matias et al. (2024) than is the lake-wide mean break-up. This suggests that the behavioral threshold identified here is more closely associated with localized deterioration of crossable ice in specific corridors than with lake-wide mean break-up, helping explain why crossing behavior shifts even when average lake-wide ice conditions remain relatively intact. In contrast, a small number of anomalous years (e.g., 2016) exhibit more spatially extensive early melt conditions, in which earlier threshold crossing occurs across a broader set of lake regions compared to typical years, resulting in an overall shift toward earlier timing at the distributional level. These years represent atypical conditions in which early loss of lake permeability is no longer confined to a limited set of localized corridors, but instead affects a larger fraction of the lake, potentially compressing or altering the migratory window relative to typical years.

Beyond this spatial heterogeneity, our system-level analysis at Contwoyto Lake indicates that, although weak monotonic trends toward earlier threshold timing and lower albedo values are statistically detectable, the availability of this crossing window appears to be shaped more by pronounced interannual variability than a smooth directional shift (Appendix S6). Although long-term climatic records document a broad trend toward earlier ice break-up across the Arctic (Magnuson et al. 2000), our 17-year lake-scale analysis shows that, over the duration of this study, and at the temporal scale experienced by individual caribou, directional warming signals are often outweighed by large year-to-year fluctuations. The behavioral consequences of this volatility are further amplified by the near-perpendicular intersection between spring migration routes and the lake, such that modest shifts in ice conditions can translate into discrete behavioral outcomes: either a direct crossing or a substantial circumnavigation. As a result, small differences in the timing or spatial extent of crossable ice surfaces can produce substantial effects on movement decisions, reinforcing the primacy of interannual variability over smooth directional trends at the scale relevant to migration. Against this backdrop of climatic volatility, the albedo thresholds identified here offer a quantitative link between environmental conditions and migratory behavior, providing a framework for linking locally realized ice conditions to migratory decision-making in dynamic permeability landscapes under climate change.

Repeated adjustments in timing or route in response to such volatility may reflect behavioral flexibility, while persistent increases in circumnavigation could reflect constraints. Such circumnavigation, especially when combined with adverse weather (Collins and Smith 1991; Eira et al. 2013), may hinder pregnant females’ timely arrival at calving grounds (Fancy and White 1987; Duquette 1988; Leclerc et al. 2021), exacerbate calving asynchrony, and reduce calf survival (Couriot et al. 2023). Monitoring changes in water-crossing behavior, especially as it relates to ice thresholds, could therefore provide an early behavioral indicator of emerging movement constraints before demographic consequences manifest (Adamczewski et al. 2022).

Beyond caribou, many cold-adapted species rely on ephemeral ice corridors to navigate fragmented northern landscapes. Our framework, though developed for caribou, is broadly applicable to migratory animals encountering dynamic surface conditions. Across the Arctic, species such as moose, muskoxen, Arctic foxes, and polar bears respond strongly to the timing and stability of ice (Pagano et al. 2021; Banfield 1954b). Globally, frozen waterways serve as critical seasonal corridors or calving refuges for Eurasian wolves (Veenbrink et al. 2026), Mongolian gazelles (Dejid et al. 2022), and Tibetan antelopes (Cao et al. 2022), underscoring that ice is not merely a constraint, but a vital and temporally bounded resource embedded in migration strategies and life histories.

More broadly, our findings demonstrate how temporally dynamic landscape features, such as ice, function as transient connectivity windows that modulate movement decisions across species (Zeigler and Fagan 2014). In such systems, plastic behavioral responses (Xu et al. 2021), and collective navigation mechanisms (Berdahl et al. 2018) often mediate interactions with shifting environmental barriers. Our approach integrates remotely sensed ice heterogeneity with individual movement trajectories to offer a transferable tool for detecting when and where functional connectivity is lost under climate pressure. Together, these insights highlight the potential of using behavioral thresholds as cross-species indicators of climate vulnerability.

By identifying a threshold at which landscape accessibility shifts rapidly, our study provides a predictive framework for understanding how climate-driven changes in permeability can reshape animal movement, with potential consequences for connectivity (Kubelka et al. 2022). Identifying quantitative thresholds, such as the albedo level at which ice becomes crossable, not only improves forecasting of behavior under changing conditions but also provides a basis for evaluating whether changing conditions are approaching limits to behavioral flexibility (Winkler et al. 2014; Eggeman et al. 2016; Courtemanch et al. 2017; Berg et al. 2019; Davidson et al. 2020; Xu et al. 2021). Ice no longer functions merely as a static barrier, but as a seasonal behavioral filter that alters landscape accessibility and, in turn, influences functional connectivity (Zeigler and Fagan 2014; Uroy et al. 2021). As migration corridors fracture under climate pressure, shifts in movement, such as threshold-based crossings and increasing circumnavigations, may provide early warning signals of emerging constraints on migratory routes. This perspective is particularly important for conservation planning (Kauffman et al. 2021), where the protection of migration corridors must account for both static structural features and dynamic environmental conditions (Reynolds et al. 2017; Moore and Schindler 2022). The integration of spatially explicit behavioral models with long-term remote sensing offers promising tools for anticipating future movement bottlenecks, guiding adaptive monitoring, and informing cross-species conservation strategies in an era of accelerating climate instability (Neumann et al. 2015; Lahoz-Monfort and Magrath 2021).

By closing the gap between landscape dynamics and individual movement decisions, our framework advances a migration ecology that is not only descriptive but increasingly predictive and relevant to conservation action under global climate change.

## Acknowledgements

We gratefully acknowledge the Government of the Northwest Territories, Department of Environment and Climate Change (GNWT-ECC), for providing the caribou GPS collar data. We thank the Handling Editor and an anonymous reviewer for constructive feedback that improved the manuscript. We are grateful to Aolin Jia for his expert advice on preprocessing MODIS albedo data. We thank Ophélie Couriot for her helpful clarification regarding collar specifications and GPS fix-rate characteristics of the NWT caribou telemetry dataset. We are thankful to the members of the Fagan Lab, Marron McConnell, Stephanie Chia, Frank McBride, Gayatri Anand, Phillip Koshute, and Jeff Demers, for their constructive input on refining the methodology. We also thank Jan Adamczewski for his valuable feedback and expert suggestions during the manuscript review. QL extends her sincere appreciation to her committee members, Sinead Farrell, Gerald S. Wilkinson, Matt Fitzpatrick, and Taylor Matthew Oshan, for their insightful guidance throughout this research. She is also thankful for the supportive academic community within the Department of Biology, the Marine, Estuarine, and Environmental Sciences (MEES) program, and the Biological Sciences Graduate Program (BISI) at the University of Maryland, College Park.

## Animal Care Protocols

All animal handling and data collection procedures were conducted in accordance with ethical guidelines and approved by the appropriate institutional animal care and use committees. The caribou collaring data used in this study were collected under permits issued by the Government of the Northwest Territories, Department of Environment and Climate Change (GNWT-ECC).

