## Appendix S1-S6 for "Dynamic Lake Ice Conditions Shape Caribou Water-Crossing Behavior in the Arctic" (from another version of this preprint: version 2, 2026-01-22)

**Appendix S1: Section 1. Different Definitions of Freeze-up and Break-up Dates**

Various definitions of freeze-up and break-up dates from in-situ data exist across the literature. For example, observations from the Swedish climate station Eklund (1999) define the "freeze date" as the first date after which the whole lake, except for small segments, remains ice-covered for at least 3 consecutive days. The "break-up date" is the day when the entire lake becomes free of ice, except for small parts of ice close to the shore or around obstacles. Conversely, Magnuson (2000) set the freeze date as the latest observed freeze date for that water body, and the breakup date as the earliest observed breakup date for that water body.

**Appendix S1: Section 2. Overview of Remote Sensing Technologies Used to Detect Lake and River Ice Phenology**

There are multiple remote sensing technologies that have been used to measure the ice phenology in lakes and rivers. Optical sensors, such as the Advanced Very High-Resolution Radiometer (AVHRR), provide daily updates with a spatial resolution of 1000 meters. However, the relatively coarse resolution of AVHRR limits its utility for smaller lakes. The Visible Infrared Imaging Radiometer Suite (VIIRS) offers improved spatial resolution at 500 meters and also provides daily updates, but its temporal coverage is shorter, as it has only been available since its 2011 launch. Recent data fusion efforts, such as Giroux-Bougard (2023), have used the OPEN-ICE algorithm to combine Sentinel-2 and Landsat 7/8 imagery at 30-meter resolution to estimate lake ice breakup dates. However, these approaches are constrained by limited temporal coverage (post-2013) due to the launch dates of the contributing satellites and are less suited for studies requiring continuous time series before that period. Similarly, the European Space Agency's Lakes Climate Change Initiative (Crétaux et al. 2022) provides MODIS-derived daily lake ice products spanning 2000–2020, but the spatial resolution is relatively coarse (1,000 meters), limiting their ability to capture within-lake heterogeneity. The

Copernicus Global Land Service (CGLS) (2022) Lake Ice Extent (LIE) product derived from Sentinel-3 SLSTR imagery offers daily 500-meter classifications across 13,000 lakes, but only since 2021. Higher resolution sensors such as the Landsat (30-meter resolution) and the Harmonized Landsat Sentinel-2 (HLS) (2–3 day temporal resolution) sensors provide snow-related indices, such as the Normalized Difference Snow Index (NDSI) (Hall and Riggs 2011). For example, Leblond (2016) used MODIS-derived normalized difference snow index (NDSI) data, averaged over 8-day intervals at a 500-meter resolution, to estimate the proportion of ice to open water across entire lakes, with values ranging from −1.0 (open water) to 1.0 (completely frozen). However, the longer temporal gaps (8 days for Landsat and 2-3 days for HLS) can make it challenging to detect rapid changes in ice phenology in complex, dynamic landscapes. Microwave sensors provide a weather-independent alternative. Passive microwave sensors, as in Matias (2024), measure Brightness Temperature to detect snowmelt onset with a spatial resolution of 3 kilometers. Meanwhile, active microwave sensors, including Synthetic-aperture radar (SAR) imaging systems, offer higher spatial resolution and are unaffected by cloud cover. Du (2015) used the SAR backscatter threshold and machine-learning algorithms to distinguish frozen and unfrozen states, while Stonevicius (2022) employed Sentinel-1 SAR data (VV and VH backscatter) to classify ice versus water in river sections. However, SAR’s low temporal resolution (i.e., revisits often exceeding five days) yields significant uncertainties when attempting to detect rapidly changing ice conditions, such as ice breakup events. SAR backscatter based on difference roughness and dielectric properties has been used to detect water and ice (Stonevicius et al. 2022), but may have difficulties distinguishing among newly formed ice, lake ice covered by wet snow, and open water (Surdu et al. 2015). Emerging technologies, including the Surface Water and Ocean Topography (SWOT) mission (launched in 2022) and NASA SnowEx data (launched in 2017), provide additional options for detecting ice melting and freezing, but only for recent years.

Despite these advancements, Moderate Resolution Imaging Spectroradiometer (MODIS) data remain one of the most practical choices for large lake systems due to its balance of spatial resolution (500 meters), temporal resolution (daily updates), and long-term data availability (launches in 1999 and 2002). For example, Pavelsky and Smith (2004) used MODIS imagery to identify ice-breakup dates for the Yenisey River in the Arctic, observing the ice/water boundaries. Cooley and Pavelsky (2016) utilized MODIS reflectance imagery to detect the ice-break-up date in the Mackenzie River in Arctic areas by dividing the river into 10 km segments and classifying the average reflectance of each river pixel in each segment as either snow, ice, or a combination. Šmejkalová (2016) used MODIS daily surface reflectance to map the ice phenology of lakes across the circumpolar Arctic, providing freeze-thaw timing datasets with both a fine temporal scale and a large spatial scale. However, the dataset in Šmejkalová (2016) is lake-point based. That is, every lake, regardless of size or location, is ultimately depicted by a single point, which is obtained from the land surface reflectance at the lake's centroid. This reduces each lake's data to a single point, overlooking within-lake spatial variation that may affect animal movement. Moreover, relying only on Terra MODIS limited her ability to obtain finer temporal resolution data.

**Appendix S1: Section 3. MODIS Land Cover Classification Background**

The MODIS Land Cover Type (MCD12Q1) Version 6.1 dataset uses supervised classifications of MODIS Terra and Aqua reflectance data to generate annual global land cover maps from 2001 to the present. And the IGBP-DIS landcover classification scheme remains extensively utilized across research fields to monitor and assess land cover changes and plays a crucial role in studies focusing on deforestation, urbanization, and climate change impacts (Song 2023; Grekousis et al. 2015).

**Appendix S1: Section 4. Rationale for Using MODIS-Derived Pure Water Pixels**

To estimate ice melt and freeze-up timing using MODIS albedo data, we relied on a lake delineation method based on MODIS-derived “pure water” pixels from the MCD12Q1.061 land cover product. MODIS is currently the only satellite product that offers daily, long-term albedo observations spanning over two decades, allowing us to align ice phenology data with caribou movement derived from GPS tracking.

Due to the 500 m spatial resolution of MODIS, a single lake is sometimes represented as multiple smaller, disjointed water patches. Therefore, we adopted a multi-lake approach based on pure water pixels rather than relying on administrative boundaries. These pixels are calculated directly from surface reflectance, enabling a more consistent and phenologically meaningful delineation of lake surfaces across time.

**Appendix S1: Section 5. Spectral Rationale for Using Albedo to Detect Ice Conditions**

Land surface albedo, defined as the ratio of the reflected shortwave radiation to the incoming solar radiation, serves as a key indicator of surface conditions (Lucht, Schaaf, and Strahler 2000). The higher the albedo, the more the surface reflects light. In the visible wavelength range (380–760 nm), snow albedo increases to 0.9–0.98, being the most reflective surface on Earth. By comparison, Arctic vegetation has a broadband albedo around 0.12 (0.06 in the visible) (Loranty, Goetz, and Beck 2011), bare ground around 0.25 (0.15), and water bodies around 0.03 (0.06) (Varotsos et al. 2014). As snow melts, the albedo decreases from 0.9 to 0.1, reflecting the transition from snow to water, absorbing the shortwave radiation as the most significant energy source (Male and Granger 1981). This transition is detectable using spectral indices in visible and shortwave infrared (SWIR) bands (Stonevicius et al. 2022).

**Appendix S1: Section 6. Kalman Filter and Climatology-Based Gap-Filling for Albedo Data**

To address gaps in daily MODIS albedo observations (due to cloud cover, sensor error, or orbital gaps), we implemented a Kalman filter-based interpolation framework, referencing daily albedo climatology from surrounding clear-sky pixels (Jia et al. 2021; 2023; Welch and Bishop 1995)

Climatology corrections depended on the number of clear-sky pixels within a clear-sky window on a given day. If there were at least three adjacent pixels within the window that provided a clear-sky albedo on that day, we added the median of the difference between the climatology value and observed values in the surrounding clear-sky pixels to the climatology value of the central pixel. We then added the resulting value to the central NA pixel on that day. If there were fewer than 3 clear-sky pixels within the clear-sky window on a given day, we used the Kalman filter based on the climatology to perform temporal interpolation on each pixel. If the pixel's initially observed albedo was NA on the first observation date, we assigned a value from the pixel's climatology value on that day.

The dynamic model with the Kalman filter only used climatology data as a changing standard. Compared to the albedo value in the previous day, the predicted albedo in that pixel kept the same degree of change (increase or decrease) as the climatology. During this, the weights of the error of climatology accumulated from the initial value and continued to increase. Once an observed value appeared, the Kalman gain was corrected by combining the weight of the observing error and the weight of the model error from climatology. We filled the NA pixel with a weighted average of the previous observation and the model-predicted value from climatology.

**Appendix S1: Section 7. Albedo Climatology**

Albedo climatology is the continuous annual series, which reflects the average of albedo observation across multiple years for each day of the year, to understand the characteristics of the specific surface and how they change over time in each pixel at the climatological scale (Jia et al. 2023). It can be used as prior knowledge to constrain the predicted albedo values of missing pixels by filtering discontinuous clear-sky data series. Combining the climatological information and filtering real-time retrievals of clear-sky days, the data gaps could be accurately predicted (Liu et al. 2013).

**Appendix S1: Section 8. The Kalman Filter**

The Kalman filter (KF) is a mathematical recursive algorithm that uses the model of how the variables are expected to change and a series of measurements observed over time to estimate unknown variables and the state of a system and to make predictions about future states (Kalman 1960). It is used in data fusion for assimilating discontinuous observations into a dynamic model (Welch and Bishop 1995).

The Kalman filter estimates a process by using a form of feedback control: the filter estimates the process state at some time and then obtains feedback in the form of (noisy) measurements. As such, the equations for the Kalman filter fall into two groups: time update equations and measurement update equations. The time update equations are responsible for projecting forward (in time) the current state and error covariance estimates to obtain the a priori estimates for the next time step. The measurement update equations are responsible for the feedback—i.e. for incorporating a new measurement into the a priori estimate to obtain an improved a posteriori estimate.

The time update equations can also be thought of as predictor equations, while the measurement update equations can be thought of as corrector equations. Indeed, the final estimation algorithm resembles that of a predictor-corrector algorithm for solving numerical problems (Welch and Bishop 1995).

In our study, the dynamic model will be established based on the albedo climatology from 2000 to 2020. Once a new observation was made available, the model prediction was updated using a weighted average of the available observations and initial modeling results. As the model iteratively predicts from the revised values, the prediction of future cloudy days will be recovered with higher accuracy. Considering the width of our research objective: lakes and rivers, the width could be very narrow, with few pixels in 500-meter resolution. 3D-KF method with a spatial window (Zhang et al. 2013; Jia et al. 2022) is not appropriate here, and we only use basic KF, which is typically utilized along the temporal dimension, to establish the temporal module.

**The Original Albedo Without Pre-processing on Contwoyto Lake in 2019**

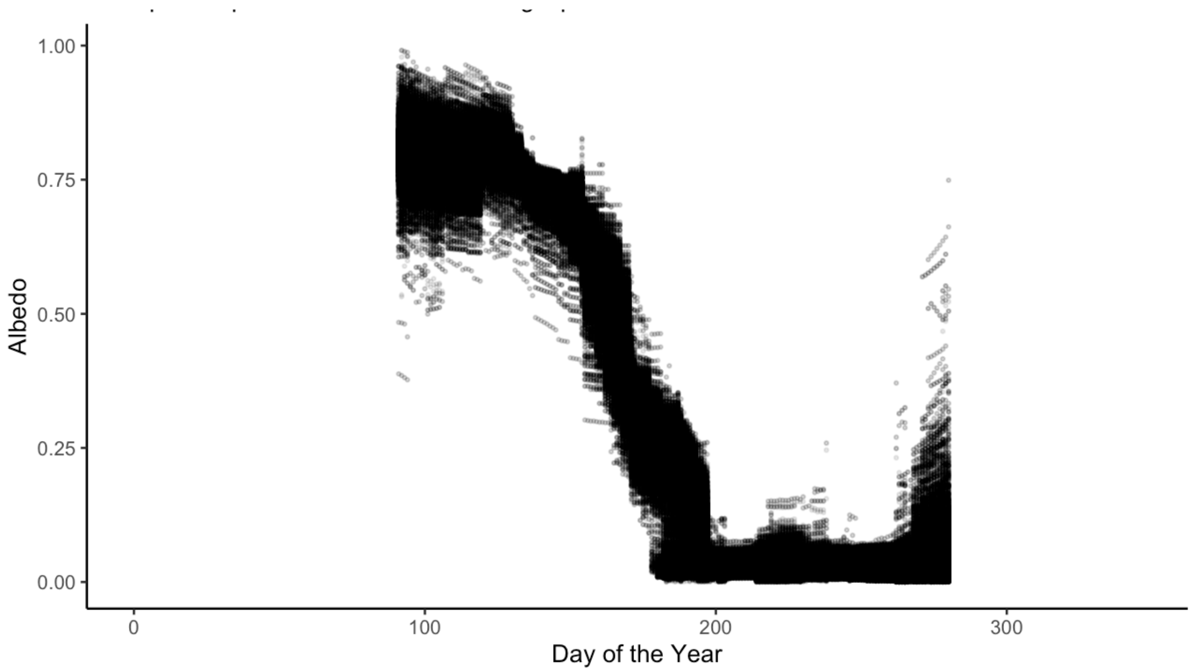


***Figure S1.*** *Albedo data from Contwoyto Lake in 2019. Dots represent the pixel-wise albedo values, with the x-axis representing the day of the year, ranging from the 92nd day (April 2nd) to the 280th (October 7th).*

### Dynamic Lake Ice Conditions Shape Caribou Water-Crossing Behavior in the Arctic

Qianru Liao, Eliezer Gurarie, and William F. Fagan

Appendix S2

**Appendix S2: Section 1. Details of the Classification of the Transit Events**

To evaluate how ice conditions affect caribou water-crossing behavior, we first determined whether a caribou likely crossed a lake or likely went around it (based on the spatial location of GPS tracking points). We classified caribou movement paths with respect to Contwoyto Lake into 3 types of transit events: "Crossing Events", "Circumnavigating Events", and "Unknown Events".

"Crossing Events" refer to cases in which one or more GPS sampling points from an individual's track actually fell within the lake surface boundaries. In such cases, it is reasonable to assume that the individual definitely crossed the lake, either by walking on ice or swimming, depending on surface conditions.

"Circumnavigating Events" represent instances where an individual's track falls within the general vicinity of the lake (intersect with 20km of the lake boundary) and intersects with the median line of the lake's long axis, but the line segments connecting consecutive GPS sampling points do not cross the lake boundaries. We applied a 20 km buffer around the lake to capture movements that approached and paralleled the lake margin without entering it, ensuring that near-lake detours plausibly reflecting avoidance of crossing were included. Sensitivity tests with larger thresholds (25 km and 30 km) yielded nearly identical results and did not identify additional Circumnavigating Events, confirming the robustness of the 20 km criterion. Thresholds much larger would extend far beyond the lake's immediate influence and thus lack ecological relevance, as such movements are unlikely to represent circumnavigation behavior. In these cases, we inferred that the individual went around the lake without crossing it.

"Unknown Events" apply when the line segments connecting consecutive GPS sampling points intersect with the lake surface boundaries, but no GPS points occurred within it. These ambiguous cases required further probabilistic classification to assess whether the caribou crossed or circumnavigated the lake. To ensure accuracy and consistency, we manually inspected the full trajectory and its spatial relationship with lake boundaries for every event. This step served as a cross-validation to confirm event type assignments prior to subsequent classification, reducing potential misclassification due to GPS error or complex shoreline geometry. Here, lake surface boundaries are defined by “pure water” pixels derived from the MODIS Land Cover Type product (MCD12Q1.061), which reflects surface reflectance rather than administrative delineations (see “Materials and Methods” section).

**Appendix S2: Section 2. Details of the Direct Path and the Circumnavigate Path**

For each Unknown Event, we considered 2 possible paths by which the caribou could have moved from one side of Contwoyto Lake to the other. These were 1) the “Direct Path,” which was the path with the shortest distance across the lake between two consecutive GPS relocations that fell on opposite sides of the lake, and 2) the “Circumnavigate Path,” which was the path with the shortest distance between the two consecutive GPS points that did not cross any water bodies between the points.

For the Circumnavigate Path, we applied the least-cost path method based on Dijkstra’s algorithm (1959) through the *shortestPath* function in the ‘*gdistance*’ R package, allowing 16-directional movement, including ‘king’s’ and ‘knight’s’ moves (Smith et al. 2018). Despite advances in shortest path algorithms (Hart et al. 1968; Sethian 1996; Mitchell 2017), Dijkstra’s algorithm remains widely used for its simplicity and effectiveness in spatial analysis, landscape ecology, geographic information systems, and geographical genetics (Adriaensen et al. 2003; Fullman et al. 2017; Etherington 2016; Lei and Church 2010; Galpern et al. 2012). The *gdistance* R package, implementing Dijkstra’s algorithm on raster cost surfaces, is widely used for modeling movement, connectivity, and resistance-based distance estimation in ecological research (Royle et al. 2013; Sutherland 1998; Etten 2017; Green et al. 2020; Weiss et al. 2018).

In our cost surface, land pixels were assigned a movement cost of 1, and water pixels (defined by MODIS-derived "pure water") were assigned a cost of NA, thereby excluding any over-water movement in Circumnavigate Paths (See the following Appendix S2: Sections 3-5 for more technical details). To compare movement strategies, we also defined a "Reference Path", consisting of the three on-land steps before and after the "Unknown Event". Figure 2 illustrates the Direct Path, Circumnavigate Path, and Reference Path for one caribou's Unknown Event.

**Appendix S2: Section 3. Background of Least Cost Path (LCP)**

The least-cost model is a widely adopted method in landscape ecology (Etherington 2016) and for predicting animal movement paths and corridors (Douglas 1994; Richard and Armstrong 2010). By generating a habitat permeability map, a grid format is used where each cell is assigned a value representing the 'cost' for an animal to traverse that specific cell (Walker and Craighead 1997; Singleton 2002) And a least cost path will accumulate the least resistance between a predefined start and end point on the resistance surface (Ray et al. 2002; Chetkiewicz and Boyce 2009) with the path being a single pixel wide on the cost map (Adriaensen et al. 2003). The least-cost path method presents a comparatively restrictive approach for creating ecological corridors because the accuracy of this method is predominantly dependent on the detailed precision of the mapped resistance surface. But at the same time, circuit theory provides a more adaptable and flexible approach by evaluating every conceivable pathway by the use of variable cutoff thresholds.  For instance, a 100% threshold would encompass the entire landscape as a corridor, whereas a 0% threshold would not include any part of the landscape (Zhao et al. 2014). Applying the theory to the research of animal's perception, the least cost path or network models (Pinto and Keitt 2009; Carroll et al. 2012) can be used to assume that animals have complete knowledge of the landscape and identify the optimal route with regard to criteria such as length and habitat quality.

**Appendix S2: Section 4. Regarding the Use of gdistance to Estimate the Shortest Circumnavigate Paths**

In heterogeneous geographic spaces represented as grids, the least-cost path between two cells and its associated distance are computed using Dijkstra's algorithm (Dijkstra 1959). This algorithm identifies the shortest path from a given source node to every other node, effectively mimicking movement routes that consider landscape obstacles and local friction, analogous to 'as the wolf runs' as opposed to 'as the crow flies.' Calculations of distance and routes begin with raster data, which are rectangular, regular grids representing continuous geographic data. Each cell within the raster is organized into rows and columns, storing specific values. Metadata accompanies rasters to describe resolution, spatial extent, and other essential properties. Raster-based distance and route calculations rely fundamentally on graph theory. Initially, raster data are transformed into graphs by connecting cell centers, converting these centers into nodes within the graph structure. The directionality connecting cells and their neighbors varies. The *gdistance* function provides three connectivity types: 1) the von Neumann neighborhood, connecting cells orthogonally to their four immediate neighbors; 2) the Moore neighborhood, connecting cells to their eight orthogonal and diagonal neighbors, forming what is known as the king's graph (reflecting all legal chess moves by a king)—a common standard in GIS software; and 3) a 16-directional connection combining king's and knight's moves. The implementation of this 16-directional connectivity in *gdistance* was inspired by the *r.cost* function from Geographic Resources Analysis Support System GRASS software (Grass Development Team 2017), also discussed in the context of distance transformations by Smith, Goodchild, and Longley (2018). This approach enhances calculation accuracy. However, because the shortest paths calculated via this method are grid-based and constrained to pass through cell centers with a maximum of 16 directional selections, the paths will be affected by the resolution of the grid. Thus, the shortest paths generated by *gdistance* can never be perfectly straight, even in scenarios where a straight line would theoretically represent the minimal distance. This introduces a slight upward bias in the estimation of shortest paths. One of us (EG) has developed an alternative method for calculating shortest paths involving a convex hull approach that will yield a straight line under those same circumstances, thus providing an improvement in simple landscapes. We have compared results for the two methods for 20 migrating caribou and found that the average shortest path is approximately 1.5% shorter when calculated using the new method than when using *gdistance*. However, at its current state of development, the alternative convex hull method does not work effectively in complex settings such as the multi-lake landscape exemplified by the Contwoyto Lake region. Specifically, concave sections of lake boundaries and alternative rotational geometries pose particular problems for the new method that have yet to be resolved. In addition, we observed that the current implementation selects lakes to include in the analysis based solely on whether they intersect the straight line connecting two GPS points. However, some lakes that do not intersect this line can still significantly influence the true shortest circumnavigation path, depending on their spatial configuration. Thus, the challenge goes beyond resolving clockwise versus counterclockwise routing, more fundamentally, we would need a way to identify all potentially influential surrounding lakes and then evaluate all relevant combinations of lake involvement in the path calculation. This quickly becomes a combinatorial problem that exceeds the scope of the current algorithm. Given these limitations and the minimal differences identified in our comparative tests, we ultimately opted to employ the shortest path methods in *gdistance* that are already in widespread use in the literature. In the next section, we provide a more detailed comparison of *gdistance* and the new method. We also include code for the new convex hull method in the code repository associated with this paper in the event that others would like to generalize and improve upon it.

**Appendix S2: Section 5. Comparison Between Least Cost Path Computed by gdistance and a Convex Hull Approach for Calculating Shortest Circumnavigate Path**

To compare the shortest circumnavigate path calculated in the method by least-cost path from *gdistance* using Dijkstra's algorithm (1959) and the shortest path method using the convex hull approach, we randomly selected 20 transit events (10 from spring migration and 10 from fall migration). We calculate their shortest circumnavigation path between the starting and ending GPS location points in a transit event on the opposite side of the lake. We will use the terms 'gdistance method' and 'convex hull method' to represent the alternative approaches.

We evaluate the difference between the results from these two methods by dividing the difference of the shortest circumnavigate distance between the gdistance method and the convex hull method by the convex hull distance to obtain the deviation rate. We found the average deviation rate to be approximately 0.015 (or 1.5%), with the difference between the two methods ranging from 0.9% to 2.7%, with 85% of cases showing a difference of less than 2% (see Table S1). Additionally, we provide several plots comparing shortest circumnavigation paths derived from each method to visualize their differences more clearly (see Figures S1–S4). These comparisons demonstrate that the shortest circumnavigation paths calculated using the gdistance method exhibit slight deviations from those generated by the convex hull method, but these deviations are minor and do not significantly affect overall path calculations.

In addition to the gdistance framework, we note that other R packages have also promoted rerouting animal tracks around geographic barriers. For example, the *pathroutr* package (London 2021) provides functions to adjust movement paths so that they avoid polygons such as lakes or rivers, ensuring biologically realistic routes when raw GPS points would otherwise fall across water or impassable features. Although we did not directly implement *pathroutr* in this study, such approaches may be valuable for future refinements in circumnavigation path delineation.

**Table S1.** Differences in the shortest circumnavigation distances calculated by the *gdistance* method and the convex hull method for 20 randomly selected transit events, including 10 events from spring migration and 10 from fall migration. The variables Year and ID identify each individual's unique trajectory. Time_before and Time_after represent the GPS timestamps at the beginning and end of each transit event on opposite sides of the lake, respectively. Crossing_duration indicates the duration of transit, calculated as the difference between Time_before and Time_after, expressed in seconds. Result_gdistance and Result_convex represent the shortest circumnavigation distances calculated using the least-cost path approach implemented in the *gdistance* package and the convex hull method developed by EG, respectively. Deviation_rate quantifies the relative difference between the two methods and is calculated as (Result_gdistance – Result_convex) / Result_convex.

| Deviation Rate | Year | ID | Time_before | Time_after | Crossing_duration (s) | Result_gdistance (m) | Result_convex (m) |
|---|---|---|---|---|---|---|---|
| 0.00920819 | 2021 | bat253 | 2021-08-16 08:00:37 | 2021-08-18 08:01:01 | 172824 | 48214.78 | 47774.86 |
| 0.009419476 | 2019 | bat205 | 2019-07-29 02:02:00 | 2019-07-29 05:01:00 | 10740 | 48067.97 | 47619.42 |
| 0.010502046 | 2017 | bat165 | 2017-05-25 15:00:00 | 2017-05-26 15:01:00 | 86460 | 25796.46 | 25528.36 |
| 0.010549283 | 2013 | bat122 | 2013-09-18 15:00:00 | 2013-09-22 15:00:00 | 345600 | 34792.08 | 34428.88 |
| 0.012292491 | 2020 | bat244 | 2020-05-06 00:00:00 | 2020-05-06 02:00:00 | 7200 | 17770.41 | 17554.62 |
| 0.01259183 | 2012 | bat106 | 2012-05-14 15:00:00 | 2012-05-15 15:00:00 | 86400 | 93461.75 | 92299.53 |
| 0.012908228 | 2010 | bev042 | 2010-05-22 15:00:00 | 2010-05-24 03:00:00 | 129600 | 59976.95 | 59212.62 |

| | | | | | | | |
|---|---|---|---|---|---|---|---|
| 0.013447825 | 2015 | bat133 | 2015-05-17 15:00:00 | 2015-05-19 15:00:00 | 172800 | 85318.98 | 84186.85 |
| 0.013730516 | 2013 | bat114 | 2013-05-22 15:00:00 | 2013-05-23 15:00:00 | 86400 | 49752.88 | 49079 |
| 0.014438582 | 2018 | bat227 | 2018-07-06 04:00:00 | 2018-07-07 20:00:00 | 144000 | 29501 | 29081.11 |
| 0.014604147 | 2011 | bat090 | 2011-07-18 12:00:00 | 2011-07-20 00:00:00 | 129600 | 23987.88 | 23642.6 |
| 0.014680924 | 2009 | bat082 | 2009-05-09 03:00:00 | 2009-05-10 15:00:00 | 129600 | 16106.7 | 15873.66 |
| 0.01539173 | 2019 | bat232 | 2019-04-16 00:00:00 | 2019-04-17 08:00:00 | 115200 | 12558.7 | 12368.33 |
| 0.017012575 | 2020 | bat279 | 2020-07-13 08:01:00 | 2020-07-15 08:00:00 | 172740 | 48628.68 | 47815.22 |
| 0.019359184 | 2005 | bat048 | 2005-07-10 13:23:00 | 2005-07-15 14:49:00 | 437160 | 120474.8 | 118186.8 |

| | | | | | | | |
|---|---|---|---|---|---|---|---|
| 0.019468<br>026 | 201<br>8 | bat23<br>2 | 2018-07-11<br>15:00:00 | 2018-07-13<br>16:01:00 | 176460 | 23235.44 | 22791.73 |
| 0.019690<br>879 | 202<br>1 | bat29<br>2 | 2021-05-20<br>08:00:11 | 2021-05-20<br>11:00:25 | 10814 | 18307.01 | 17953.49 |
| 0.020165<br>728 | 201<br>4 | bat12<br>2 | 2014-05-11<br>15:00:00 | 2014-05-12<br>15:00:00 | 86400 | 102331.6 | 100308.8 |
| 0.020287<br>191 | 201<br>7 | bat15<br>6 | 2017-07-14<br>16:01:00 | 2017-07-18<br>08:00:00 | 316740 | 53880.03 | 52808.69 |
| 0.027575<br>455 | 202<br>0 | bat26<br>9 | 2020-07-23<br>16:00:00 | 2020-07-25<br>00:01:00 | 115260 | 18336.19 | 17844.13 |

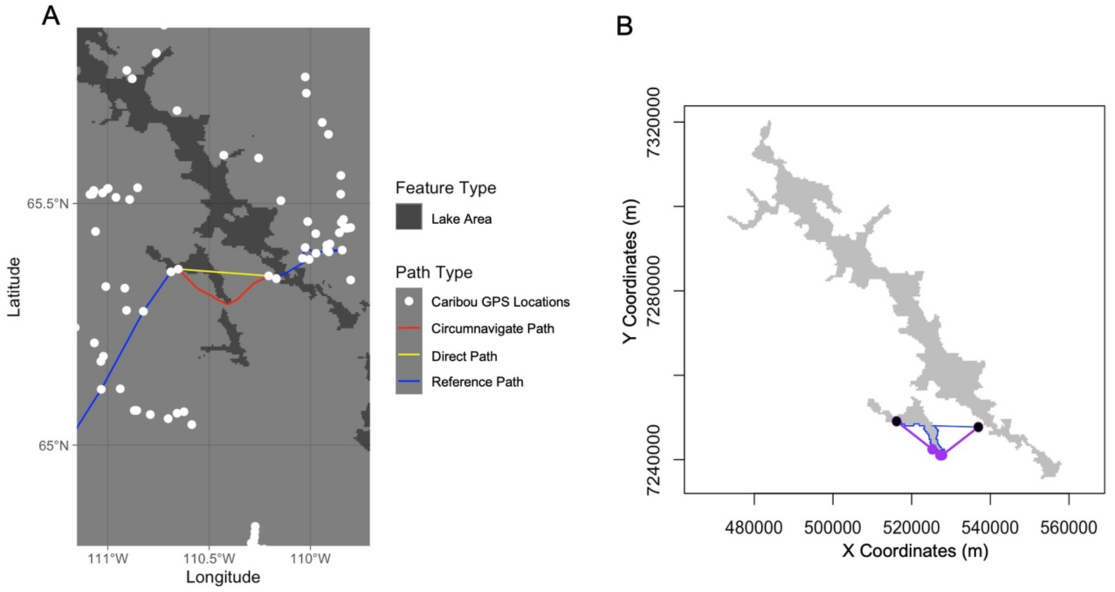


***Figure S1.*** *This figure illustrates the shortest circumnavigation paths for the transit event of individual ID 'bat165' in 2017, calculated using the gdistance method and the convex hull method. (A) The Circumnavigate Path (shortest circumnavigation route) calculated using the gdistance method is shown in red. The Direct Path is shown in yellow, the Reference Path is shown in blue, and the GPS location points along this trajectory are indicated in white. Dark regions represent the Contwoyto Lake areas. (B) The Circumnavigate Path calculated using the convex hull method is depicted in purple. Black points indicate the starting and ending locations of the transit event.*

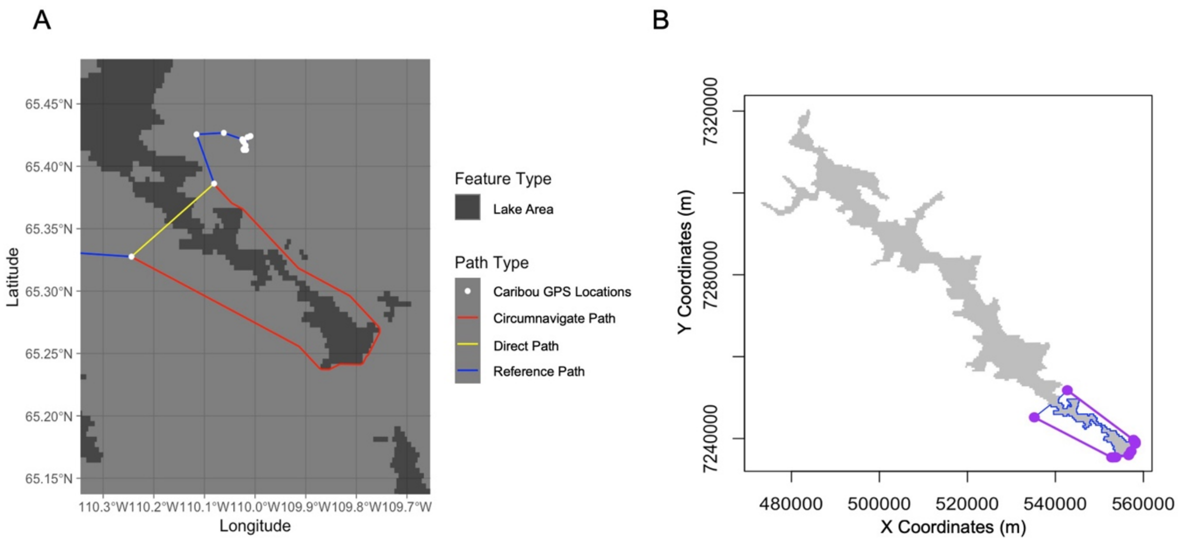


***Figure S2.*** *This figure illustrates the shortest circumnavigation paths for the transit event of individual ID 'bat205' in 2019, calculated using the gdistance method and the convex hull method. (A) The Circumnavigate Path (shortest circumnavigation route) calculated using the gdistance method, is shown in red. The Direct Path is shown in yellow, the Reference Path is shown in blue, and the GPS location points along this trajectory are indicated in white. Dark regions represent the Contwoyto Lake areas. (B) The Circumnavigate Path calculated using the convex hull method, is depicted in purple. Black points indicate the starting and ending locations of the transit event.*

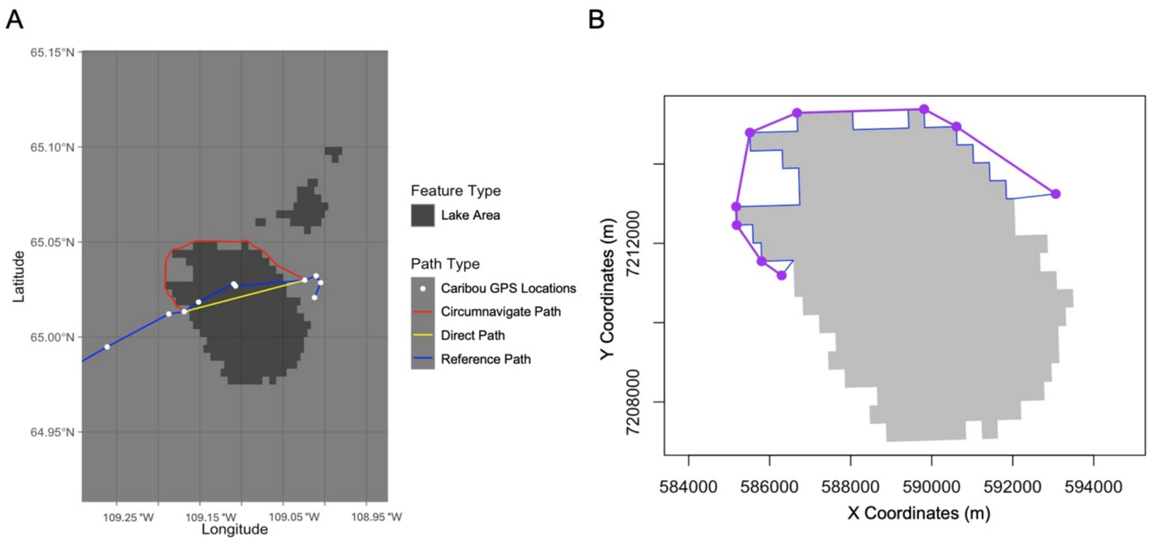


***Figure S3.*** *This figure illustrates the shortest circumnavigation paths for the transit event of individual ID 'bat232' in 2019, calculated using the gdistance method and the convex hull method. (A) The Circumnavigate Path (shortest circumnavigation route) calculated using the gdistance method is shown in red. The Direct Path is shown in yellow, the Reference Path is shown in blue, and the GPS location points along this trajectory are indicated in white. Dark regions represent the Contwoyto Lake areas. (B) The Circumnavigate Path calculated using the convex hull method is depicted in purple. Black points indicate the starting and ending locations of the transit event.*

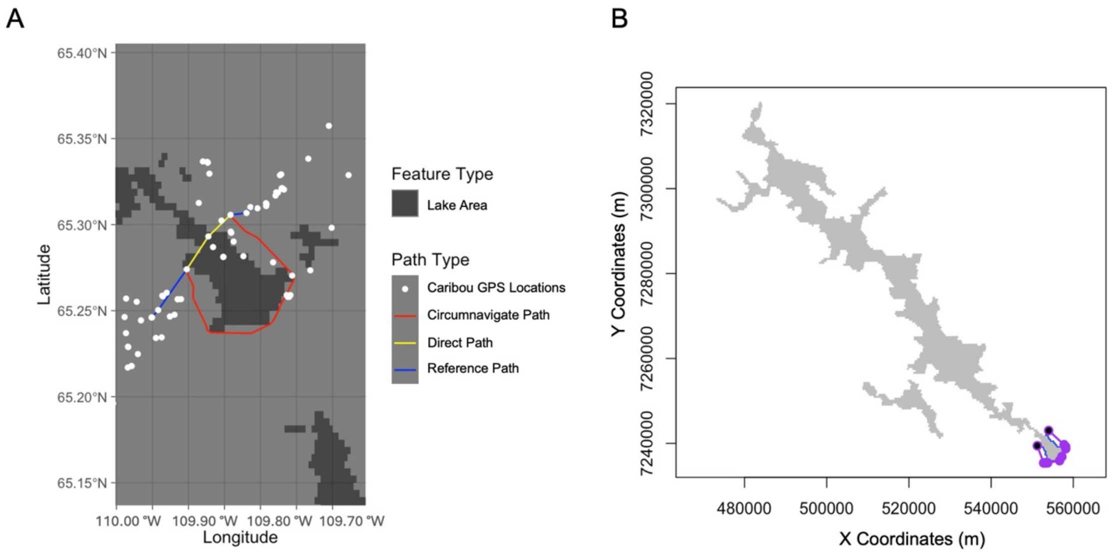

***Figure S4.*** *This figure illustrates the shortest circumnavigation paths for the transit event of individual ID 'bat244' in 2020, calculated using the gdistance method and the convex hull method. (A) The Circumnavigate Path (shortest circumnavigation route) calculated using the gdistance method is shown in red. The Direct Path is shown in yellow, the Reference Path is shown in blue, and the GPS location points along this trajectory are indicated in white. Dark regions represent the Contwoyto Lake areas. (B) The Circumnavigate Path calculated using the convex hull method is depicted in purple. Black points indicate the starting and ending locations of the transit event.*

### Dynamic Lake Ice Conditions Shape Caribou Water-Crossing Behavior in the Arctic

Qianru Liao, Eliezer Gurarie, and William F. Fagan

Appendix S3

#### Appendix S3: Section 1. Effect of GPS Fix Interval on Event Classification

GPS fix rates varied within and among individuals and years due to differences in collar programming. To evaluate how temporal resolution affects our ability to classify movement events (Crossing vs. Circumnavigating vs. Unknown), we quantified temporal resolution using the GPS fix interval associated with the reference path surrounding each transit event (Table S1), rather than the total event duration itself. Specifically, we calculated the median time interval between consecutive GPS fixes for the three steps before and the three steps after each transit event. We then grouped these intervals into bins that reflect these operational sampling resolutions (e.g., 1 h, 3h, 6 h, 12 h, 24 h, etc.) rather than arbitrary equal-width bins. The percentage of "Unknown" events increased substantially with coarser temporal resolution (from 27.3% at ≤1 h to 74.6% at 12-24 h), indicating that classification uncertainty is partly driven by GPS sampling interval rather than behavioral ambiguity.

**Table S1.** Number of caribou movement events (Crossing, Circumnavigating, and Unknown) categorized by GPS fix interval bins.

| Event-level GPS Fix Interval (hours) | Crossing Event | Circumnavigating Event | Unknown Event | Total Event | Unknown Event (% of event in bin) |
|---|---|---|---|---|---|
| **<=1** | 6 | 2 | 3 | 11 | 27.3% |
| **1-3** | 25 | 32 | 56 | 113 | 49.6% |
| **3-6** | 6 | 28 | 15 | 49 | 30.6% |
| **6-12** | 15 | 38 | 76 | 129 | 58.9% |
| **12-24** | 10 | 6 | 47 | 63 | 74.6% |
| **>=24** | 0 | 1 | 1 | 2 | 50% |
| **Total Number** | 62 | 107 | 198 | 367 | NA |

Note: GPS fix interval was defined as the median temporal interval of on-land reference steps surrounding each transit event (three steps before and after). Percentages indicate the proportion of Unknown Events within each GPS fix interval category.

**Appendix S3: Section 2. Definition of "Starting Point" and "Ending Point" for Each Transit Event**

To compare caribou transit speeds across or around Contwoyto Lake, we first identified the "Starting Point" and "Ending Point" for each event, defined as GPS locations on opposite sides of the lake.

For Crossing Events, by definition, one or more sampling points fell within the boundaries of the lake, and the "Starting Point" and "Ending Point" were on opposite sides of the lake but were not consecutive GPS relocations. For Unknown Events, the "Starting Point" and "Ending Point" were two consecutive GPS relocations that fell on opposite sides of the lake. For Circumnavigating Events, we used a 20 km buffer around the lake as a guiding framework to identify the "Starting Point" and "Ending Point," defined as the last GPS relocation before entering and the first GPS relocation after leaving the buffered lake region on the opposite side. In practice, the actual start and end points could fall within a smaller or slightly larger distance than 20 km, depending on the geometry of the circumnavigation. This flexible approach, combined with visual inspection of each trajectory, was applied to ensure accuracy—that is, to confirm that the selected events indeed represented true circumnavigation of the lake.

**Appendix S3: Section 3. Definition of Three Types of Speeds for Each Transit Event**

Speed was then calculated as the distance traveled divided by the duration of the time interval(s) between the "Starting Point" and "Ending Point" for each event.

We derived three types of speeds for each event, including 1) the "Direct Speed", which was the average speed at which the caribou would have moved between the Starting and Ending Points if it had traveled via the straight path directly across the lake, 2) the "Circumnavigate Speed", which was the average speed that the caribou would have traveled if it had gone between the Starting and Ending points via the shortest overland route, and 3) the "Reference Speed," which was the average speed of the caribou across the three tracked steps before and after the transit event.

**Appendix S3: Section 4. Definitions of Ten Predictor Variables Used to Classify Unknown Events**

1. **Speed Ratios**

We calculated three speed ratios to characterize movement dynamics during transit events: Circumnavigate Speed / Reference Speed, Direct Speed / Reference Speed, and Circumnavigate Speed / Direct Speed.

2. **Albedo Indices at Small, Medium, and Large Spatial Scales**

All ice-related predictors listed below are expressed as albedo percentile rank (APR), a within-pixel, within-year standardized index, rather than raw albedo. APR is used to normalize strong interannual variability in absolute albedo caused by snow depth, surface wetness, illumination geometry, and atmospheric effects, and should not be interpreted as a physical fraction of ice cover.

Albedo indices were calculated at three spatial scales to quantify ice conditions. At the smallest scale, we have "APR at the Nearest Pixel," which is simply the APR of the pixel that includes the Starting Point for a given transit. At an intermediate scale, we have "APR Along the Potential Crossing Path," which was calculated as the average APR along a putative transit route. Lastly, at the largest scale, we have the "APR of the Entire Lake Area," which is the average APR of all the pixels within Contwoyto Lake on the day of transit.

**3. Lake Variables**

Two variables quantified interactions between caribou and lake features. First, "Lake Width" was the straight-line width of the caribou's potential trajectory across the lake. Second, "Proportion of Time Spent in the Lake Area" was the estimated proportion of the total transit time that the individual spent within the boundaries of the lake. We estimated this time using the *occurrence* function of the *ctmm* R package by calculating the product of the crossing duration and the sum of the probability mass across all lake pixels in which the caribou was estimated to occur. The *ctmm* package is well-suited for handling our GPS movement data with irregular sampling intervals, as it builds continuous-time movement models that account for autocorrelation and explicitly address the issue of variable sampling rates (Fleming et al. 2014; 2016; 2017; Calabrese et al. 2016).

The crossing duration was calculated as the time difference between the Starting and Ending Points for each transit. Because the sampling interval and the crossing duration can vary widely among individuals, we relied on the proportion of time each individual spent within the lake boundaries.

**4. Individual Sex**

Sex was included as a variable due to behavioral differences, particularly during the spring migration, when pregnant females prioritize timely arrival at calving grounds.

**5. Migration Season**

Migration season was defined based on observed movement patterns. "Spring Migration" included all transit events occurring before July 1st, whereas "Fall Migration" included all transit events occurring after July 1st.

**Appendix S3: Section 5. Definitions of Seven Metrics for Evaluating Predictive Model Performance**

We trained both Random Forest (RF) and Binomial Logistic (BL) models using the ten predictor variables described above. Events were based on a 50% crossing probability threshold, where events with predicted probabilities >=50% were labeled as Crossing Events and those with less < 50% as Circumnavigating Events. Classification results were compared across models to assess prediction consistency and validate the robustness of classification outcomes. Additionally, both models identified the most important predictors of caribou water-crossing behavior.

We also used 80% of the dataset to train the models and reserved 20% for testing. Training data were used to build and optimize the models, while testing data assessed model predictive performance. Model performance was evaluated using the following seven metrics: Accuracy, Sensitivity, F1 Score, Kappa Value, Area under the curve (AUC), delta Akaike information criterion (delta AIC), and Out-of-Bag (OOB) error. Here are the details:

1) "Accuracy" measures the proportion of correctly classified events.

2)"Sensitivity" measures the ability of the model to correctly classify Circumnavigating Events.

3) "F1 Score" is the harmonic average of Precision and Recall, where Precision is the proportion of the true positive results among all predicted positive samples (true positives and false positive), while Recall refers to the proportion of the true positive results among all actual positive samples (true positives and false negative). The formula for calculating the F1 Score is: $F1\ Score = \frac{(2 \times Precision \times Recall)}{(Precision + Recall)}$. By combining Precision and Recall, the F1 score provides a balanced evaluation of a model's performance, especially for imbalanced class samples, and serves as a complementary metric with "Accuracy" (Powers 2011; Tharwat 2021; Bala et al. 2024).

4) “Kappa Value” is the measure of agreement between training data and test data, ranging from 0 (no agreement) to 1 (perfect agreement), providing an overall assessment of the accuracy of the classification.

5) “Area under the curve (AUC)” measures the model’s ability to discriminate between classes, ranging from 0.5 (random prediction) to 1.0 (perfect prediction).

6) “Delta Akaike information criterion (delta AIC)” evaluates the difference in AIC score between the best model and the next best model being compared for the BL model.

7) “Out-of-Bag (OOB) error” measures the prediction error of the RF model.

**Appendix S3: Section 6. Details of Addressing Data Imbalance in the RF Model**

For the RF model, we used the cost-sensitive method to address the imbalance in the dataset (e.g., fewer Known Circumnavigating Events than crossing events in spring migration and the opposite in fall migration). Class weights were assigned based on the inverse frequency of each class using the *class_wt* parameter in the *randomForest* function. This approach outperformed other methods that we explored, such as oversampling, undersampling, Random Over-Sampling Examples (ROSE), and Synthetic Minority Oversampling Technique (SMOTE), due to a better performance of the cost-sensitive method as judged by higher F1 scores and sensitivity, and lower Out-of-Bag (OOB) error.

### Dynamic Lake Ice Conditions Shape Caribou Water-Crossing Behavior in the Arctic

Qianru Liao, Eliezer Gurarie, and William F. Fagan

Appendix S4

**Appendix S4: Section 1. The Spatial Pattern of Ice Melting and Freezing Timing During Spring and Fall Migration**

To further characterize the ice dynamics, we calculated the first date of each year on which the albedo in individual pixels reaches specific percentiles between the maximum and minimum annual albedo values. This analysis allowed for a detailed analysis of ice-breaking patterns in the spring migration (Figure S1), the fall migration (Figure S2), and the whole observation period (Figure S3).

During spring migration, the ice melting patterns showed notable spatial variation. Above the 60th percentile, small clusters of lower albedo pixels (darker colors) gradually expand across the lake, with greater spatial heterogeneities as the percentile drops. When the percentile is 60%, the timing of ice melt is quite homogeneous spatially. Below the 60% percentile, the southern part of the lake exhibited earlier declines in albedo compared to the northern region.

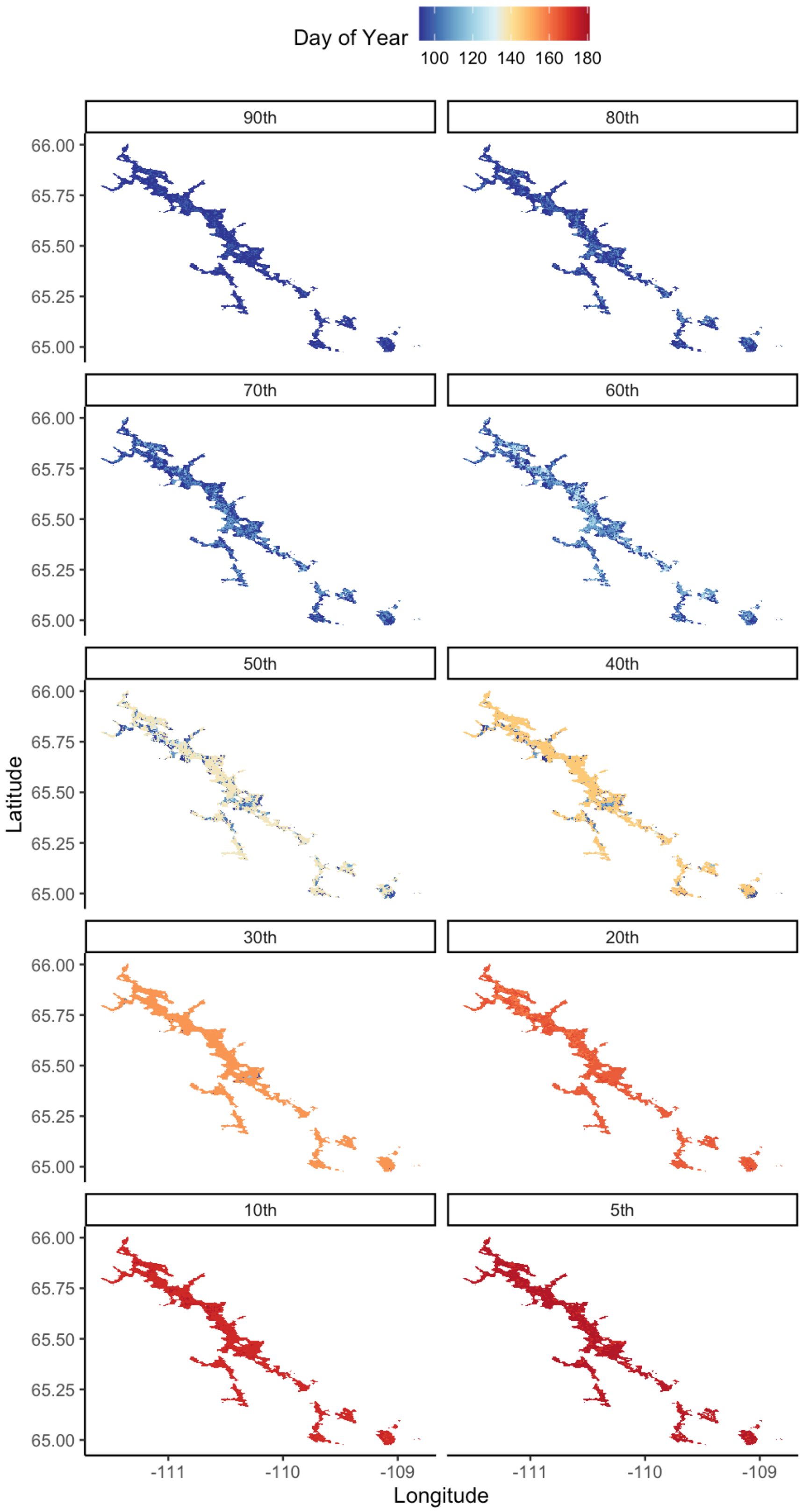


***Figure S1.*** *The first date the albedo of Contwoyto Lake pixels declined to particular percentiles (from 90th to 5th) of the maximum albedo recorded of pure water body pixels. Legend colors*

*represent the day of the year from April 2nd (92nd day, dark blue) to July 1st (182nd day, dark red) during the spring migration period.*

During the fall migration, the ice-freezing patterns of Contwoyto Lake differed significantly from the ice-melting patterns observed in spring. Below the 30th percentile, ice formation occurred earlier in the narrower sections of the lake compared to the wider areas. Above the 30th percentile, the timing of ice formation was more uniformly distributed from earlier days to later days, changing from light red to dark red.

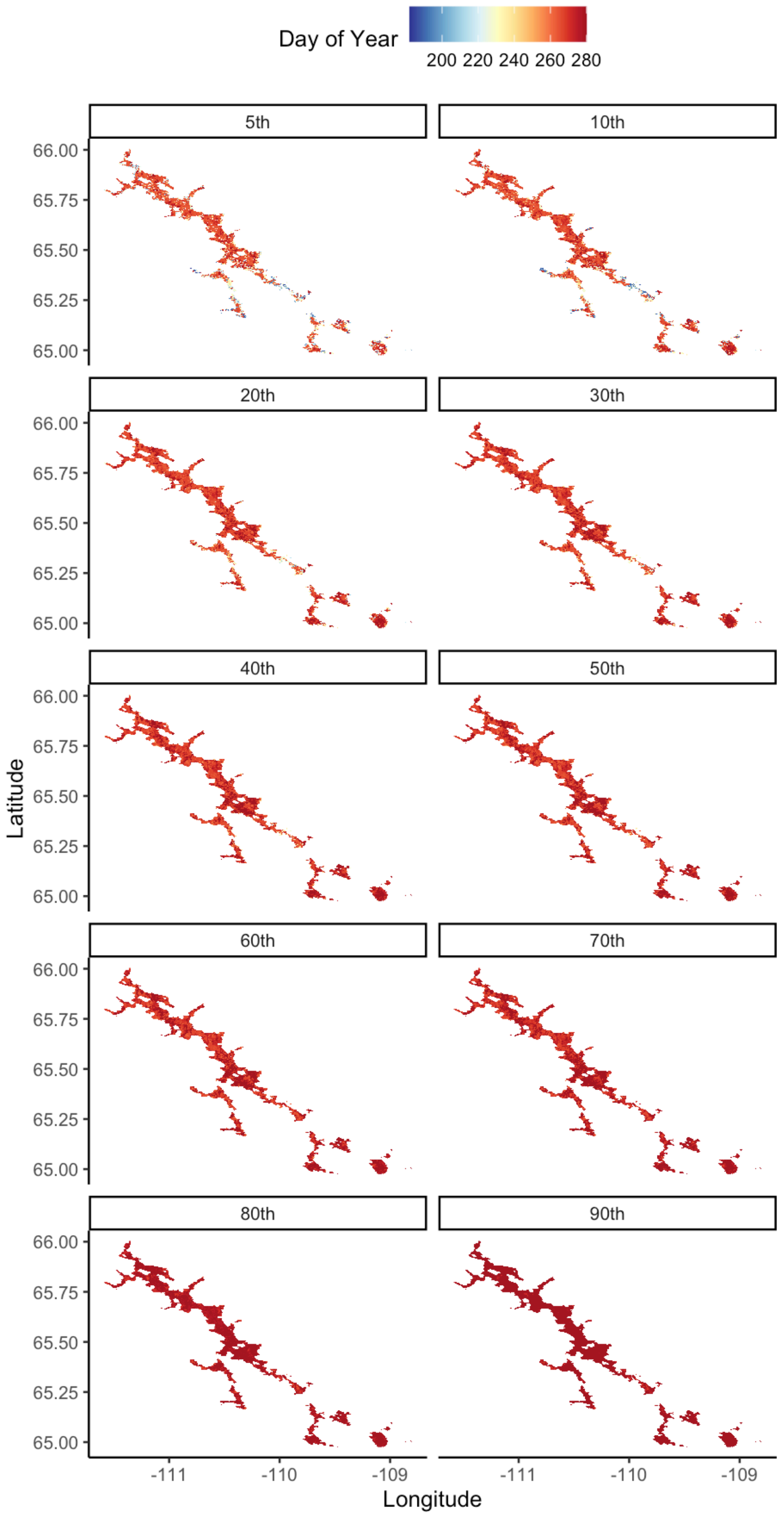

Day of Year
200 220 240 260 280
5th
10th
20th
30th
40th
50th
60th
70th
80th
90th
66.00
65.75
65.50
65.25
65.00
Latitude
-111
-110
-109
Longitude

***Figure S2.*** *For 2019, the first date the albedo of Contwoyto Lake pixels declined to particular percentiles (from 5th to 90th) of the maximum albedo recorded of pure water body pixels. Legend colors represent the day of the year from July 2nd (183rd day, dark blue) to Oct 7th (280th day, dark red) during the fall migration period.*

**Whole Observation Period**

The timing of ice melting during the whole observation period, ranging from the 92nd to the 280th day of the year (Figure S3), reveals distinct temporal patterns. The earliest date when the albedo dropped to the 90th percentile was after the 180th day (June 29th), while the earliest day when the albedo dropped to the 5th percentile was beyond the 250th day (Sep 7th). The albedo of the northern part of the lake consistently declined later than the southern part. In contrast, the timing of the freezing for the entire observation period lacks sufficient data to capture the transition from the 5th to the 90th percentile due to the polar night. The latest observation date of the year is Oct 7th, which occurs just about one month after the albedo reached its lowest value, limiting the ability to comprehensively analyze freezing patterns over the full range.

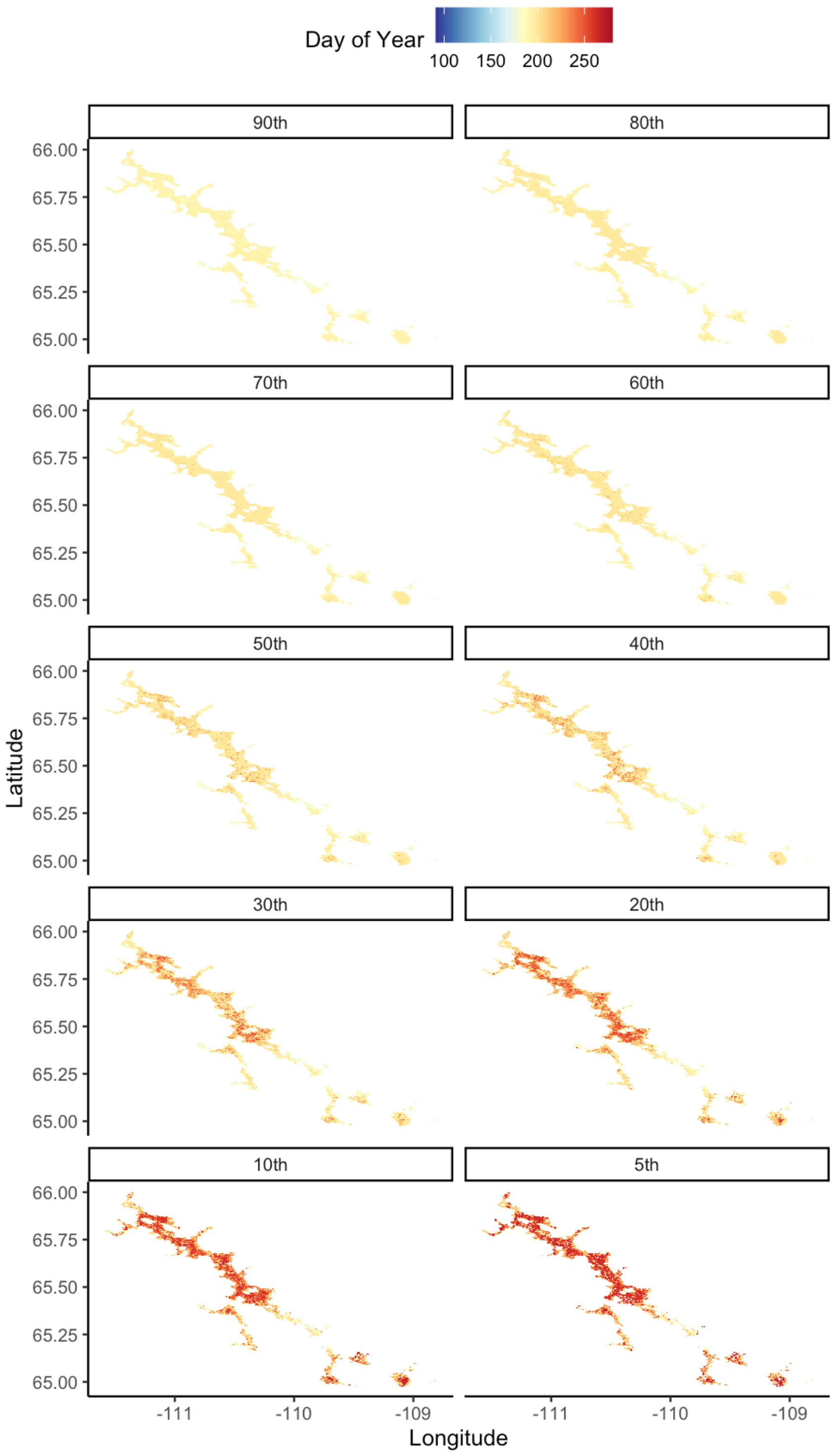


***Figure S3.*** *For 2019, the first date the albedo of Contwoyto Lake pixels declined to particular percentiles (from 90th to 5th) of the maximum albedo recorded of pure water body pixels. Legend colors represent the day of the year from April 2nd (92nd day, dark blue) to Oct 7th (280th day, dark red) during the whole observation period, excluding the polar night.*

**Appendix S4: Section 2. Spatial Distribution of Crossing Events (Within Lake Boundaries)**

A total of 109 GPS relocation points from 47 individuals occurred within the boundaries of Contwoyto Lake in the crossing events.

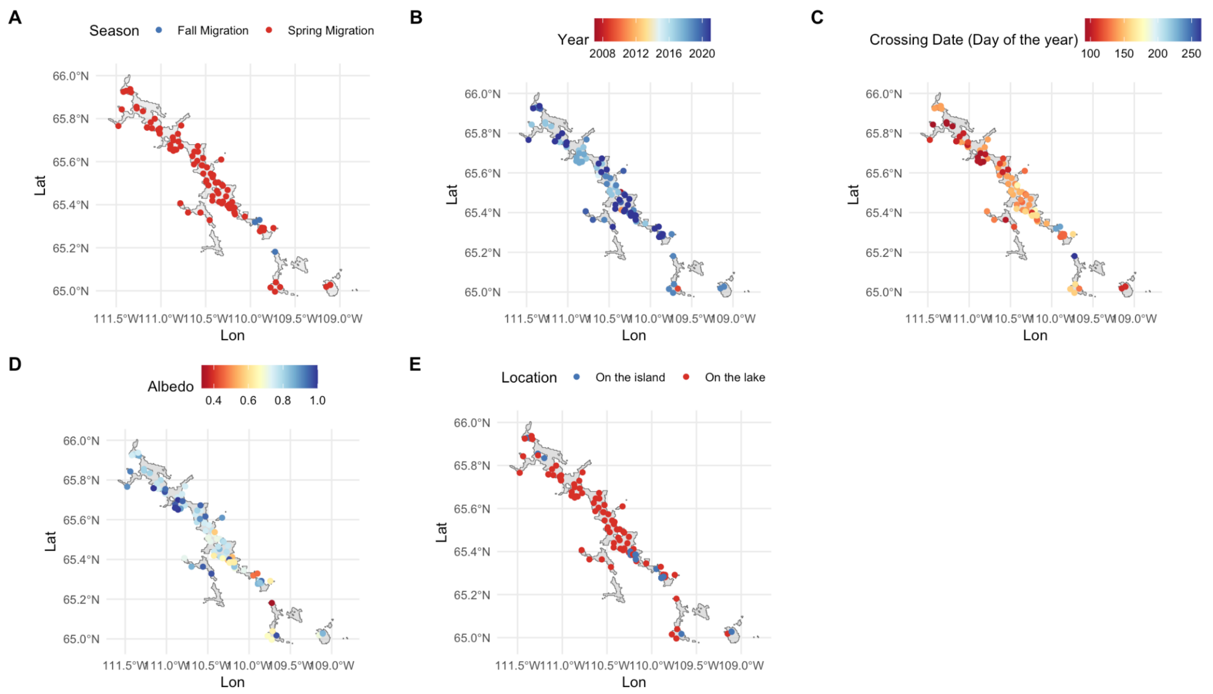


***Figure S4.*** *Overview of the spatial pattern of known crossing events from 2001 to 2021. (A) Examines the spatial distribution of known crossing events in different migration seasons: spring migration before July 1st (Red) and fall migration after July 1st (Blue). (B) Shows the spatial distribution of known crossing events in different years, from red (smaller year) to blue (larger year). (C) Depicts the spatial distribution of known crossing events that happened on different crossing dates (day of the year), from red (earlier) to blue (later). (D) Illustrates the albedo in percentile rank where the known crossing events occurred, from red (0) to blue (1). (E) represents the landscape where the known crossing occurred, on the pure lake areas (Red) or on the lake island (Blue).*

### Dynamic Lake Ice Conditions Shape Caribou Water-Crossing Behavior in the Arctic

Qianru Liao, Eliezer Gurarie, and William F. Fagan

Appendix S5

**Appendix S5: Section 1. Detailed Results and Cross-Model Validations in the "Spring Only" Dataset**

For the "Spring Only" dataset, the RF model achieved 100 % prediction accuracy with an Out-of-Bag OOB error rate of 3.51 %. The F-1 score was 1.0, and the Area Under the Curve (AUC) was also 1. These results indicate that the model successfully distinguished between the crossing and circumnavigating events.

The variable importance analysis, as shown by the mean decrease in Gini Importance Index (Figure 6A), identified the top three most important variables for classifying unknown events in the RF model: APR Along the Potential Crossing Path, Circumnavigate Speeds/Direct Speeds (log), Circumnavigate Speeds/Reference Speeds (log).

The best BL model was selected based on coefficient regularization and elimination of multicollinearity using Lasso regression, followed by evaluation of all variable combinations with the *dredge* function in R. Visual inspection of binned averages revealed monotonic relationships, with no evidence of U-shaped or inverted-U patterns. For this reason, quadratic terms were not retained in the main model, as the linear specification already captured the observed trend. This model was selected for its low ΔAIC and identified the APR along the potential crossing path (P value $< 0.05$) as the variable that most affected crossing behavior. The model achieved 95.8 % prediction accuracy, 0.96 sensitivity, an F1 score of 0.974, a ΔAIC of -0.51 (relative to the next best converged BL model), and AUC of 0.99, showing a robust classification performance. The predicted crossing probability showed a positive relationship with the APR along the potential crossing path (Figure 6B), where higher APR values correspond to increased probabilities of crossing events. The threshold for a 50% crossing probability threshold corresponded to an albedo percentile rank of 0.56.

A comparison of the predicted classification results for unknown events between the RF and BL models shows that all crossing probabilities exceed 50% in both the RF model and the BL model. This indicates high accuracy and strong agreement between two models in classifying unknown events (Figure 6B).

**Cross-model Validation**

By setting the threshold of crossing probability as 0.5, events with a probability higher than 0.5 were classified as crossing (1), while those with a probability of 0.5 or lower were classified as circumnavigating (0) in the confusion matrix (Table 2). The results revealed 100% consistency between the predictions of the two models. Both RF and BL models classified all 73 unknown events as crossing events.

**Appendix S5: Section 2. Detailed Results and Cross-Model Validations of the "Fall Only" Dataset**

Among the 97 known events during the fall migration, we found only 3 crossing cases compared to 94 circumnavigating cases, resulting in an extremely unbalanced dataset. This imbalance significantly affected the performance of the RF model, which achieved a 0% OOB estimate of error rate, but struggled to predict crossing events accurately. The best performance model had 95% accuracy in predicting the test data but had 0% sensitivity and an F1 score of 0. With only 2 crossing events in the training data, the model failed to predict the single crossing event in the test set, indicating instability in correctly identifying crossing behavior.

The top 3 most important predictive variables in the RF model in the fall migration were the Direct Speeds/Reference Speeds (log), Circumnavigate Speeds/Reference Speeds (log), and Proportion of Time Spent in Lake Area. Notably, none of these variables were related to albedo (Figure S1A). With low APR (<0.5) in fall migration, most caribou crossing events likely involved swimming rather than crossing on ice-covered lake surfaces.

In the BL model, we applied Lasso and Ridge regression to reduce multicollinearity and identify the most influential variables. After evaluating all variable combinations using the *dredge* function in R (Bartoń 2024), the best model was identified based on its performance and lowest ΔAIC. Visual inspection of binned averages revealed monotonic relationships, with no evidence of U-shaped or inverted-U patterns. For this reason, quadratic terms were not retained in the main model, as the linear specification already captured the observed trend. Specifically, for the APR of the Entire Lake Area in the fall-only dataset, a few bins with small sample sizes produced minor fluctuations in crossing probability ($p = 0.1–0.2$ in two bins with $n = 10$ each). This led smooth GAM fits to appear statistically significant (LRT $p < 0.001$), but the apparent nonlinearity reflects overfitting to sparse data rather than a meaningful biological process. We therefore retained the linear specification, consistent with the interpretation that low albedo values correspond primarily to swimming events rather than over-ice crossings.

The model that identified Circumnavigate Speeds/Reference Speeds (log) (P value < 0.01) as the variable that most affected crossing behavior, had high predictive performance, with an accuracy of 97.94%, a sensitivity and F1 value of 0.667, a kappa value of 0.656, a ΔAIC of -0.75 (compared to the next best converged BL model) and a high AUC of 0.99. The threshold for the log ratio of the circumnavigate speed over reference speed was approximately 2.698 (Figure S1B).

**Cross-model Validation**

To evaluate the predictive performance of the models in classifying unknown events, we compared the crossing probabilities obtained from the RF model and the BL model. A 50% crossing probability threshold was applied to classify events as either "crossing" or "circumnavigating". And the consistency of classifications between the RF and BL models was then assessed by comparing the predicted results using a confusion matrix.

We found that 122 unknown events were consistently classified as circumnavigating by both the RF and BL models. One event was predicted as circumnavigating by the BL model but as crossing by the RF model, and another showed the opposite pattern. Only one unknown event was classified as crossing by both models (Figure S2). Overall, model predictions were

98.40% consistent, underscoring the importance of the speed ratio in distinguishing between crossing and circumnavigating events during fall migration.

**Appendix S5: Section 3. Detailed Results and Cross-model Validations in "Combined Dataset**

Unlike the individual spring and fall datasets, which are unbalanced in opposite directions, the spring dataset contains a large number of crossing events and few circumnavigating events, while the fall dataset contains many circumnavigating events and few crossing events. The combined dataset results in a more balanced distribution of crossing and circumnavigating cases. Although caribou exhibit distinct ecological movement patterns between seasons, with spring migration characterized by more directed long distance migrating movement and fall migration showing more nomadic behavior, we chose to use the combined dataset to identify variables that have an overall influence on water crossing behavior across all events.

The RF model applied to the combined dataset, including both spring and fall migration, demonstrated strong classification performance. The model achieved a prediction accuracy of 91.18% on the test data, with an F1 score of 0.90, sensitivity of 0.87, and a kappa value of 0.82. The Out-of-Bag (OOB) error rate was 1.48%, underscoring the model's reliability. Additionally, the Area Under the Curve (AUC) was 0.98, indicating excellent ability to distinguish between crossing and circumnavigating events. According to the Gini Importance Index, the top three most important variables were the APR along the potential crossing path (Medium Scale), the APR at the nearest pixel, and the APR at the scale of the entire lake (Figure S1C).

The best performing BL model identified three key predictors: APR Along the Potential Crossing Path (P value ~ 0), Proportion of Time Spent in Lake Area (P value < 0.05), and Circumnavigate Speeds/Reference Speeds (log) (P value < 0.05). All three variables were significant and positively associated with crossing probability (Table 2). These predictors were selected following coefficient regularization with Lasso Regression to remove multicollinearity, and the final model was chosen based on the evaluation of all variable combinations using the *dredge* function in R. Visual inspection of binned averages revealed monotonic relationships, with no evidence of U-shaped or inverted-U patterns. For this reason, quadratic terms were not retained in the main model, as the linear specification already captured the observed trend.

We also evaluated models that included migration season as a main effect and in interaction with albedo or lake-use time. Although these specifications achieved lower AIC values (16.4–22.1), they produced warnings of quasi-complete separation, with extreme coefficient estimates and predicted probabilities collapsing to 0 or 1. This outcome indicates that season, when combined with environmental predictors, nearly perfectly classified events and destabilized parameter estimation. By contrast, the linear specification without season (AIC = 23.3) provided stable and interpretable coefficients, and was therefore selected as the preferred model. This best-fitting BL model achieved a prediction accuracy of 98.8%, ΔAIC of –2.12 (compared to the next best-converged BL model), a sensitivity of 0.98, a kappa value of 0.97, and an AUC of 0.99.

Although our best-performing BL model did not retain season as an explicit predictor, the results nonetheless highlight the strong role of seasonal context. The influence of albedo on crossing behavior differs markedly between spring and fall: when albedo is low (less than 0.5), observed crossings correspond to swimming events rather than crossing movements over ice (Figure S1D). This pattern aligns with the distinct ecological patterns of spring and fall migrations—spring being more directed and long-distance over ice, fall more nomadic around the large open-water bodies—and underscores that season strongly shapes the mechanisms by which environmental conditions affect water-crossing decisions. By using the combined dataset, our goal was not to mask these differences, but to identify predictors that consistently influence crossing probability across both seasons.

Comparing the most important variables across the spring, fall, and combined datasets reveals a consistent pattern. APR along the potential crossing path (identified as the most important variable in the spring dataset) and the log ratio of the circumnavigate speed over reference speed (identified as the most important variable in the fall dataset) remain two of the top predictors of crossing behavior in the combined dataset. In addition, the proportion of time spent within lake areas also emerged as a key variable in the combined model.

These findings highlight the critical role of ice conditions, reflected by albedo, in influencing crossing behavior and demonstrate the consistency of key predictors in capturing the underlying mechanisms that shape caribou decision-making during migrations.

**Cross-model Validation**

To evaluate the prediction performance of the models in classifying unknown events, we compared the crossing probabilities predicted from the RF model and BL model and set 50% as the threshold of crossing. The predicted probabilities for the entire dataset, which combined both spring and fall migration seasons, showed no statistically significant difference in prediction results between the RF and BL models based on a t-test (P value = 3.1e-16).

We also set the threshold of crossing probability as 0.5 and classified events with a probability higher than 0.5 as crossing (1) and lower than 0.5 as circumnavigating (0) in the confusion matrix. We found that 67 unknown events were consistently predicted as circumnavigating events by both the RF and BL models. Additionally, 58 unknown events were predicted as crossing events in the BL model and as circumnavigating events in the RF model. A total of 73 unknown events were classified as crossing events by both models. The agreement between the RF and BL models was 70.71%, indicating that the models demonstrated a higher-than-average level of consistency in classifying unknown events.

Both the Random Forest (RF) and Binomial Logistic (BL) models demonstrated strong classification performance in distinguishing crossing and circumnavigating events. Across all datasets (spring, fall, and the combined dataset), the consistency in predictions between the two models exceeded 70%, and the accuracy of each model surpassed 90%. These results highlight the robustness of the classification framework and support the reliability of key predictors derived from remotely sensed lake albedo and other movement-related variables.

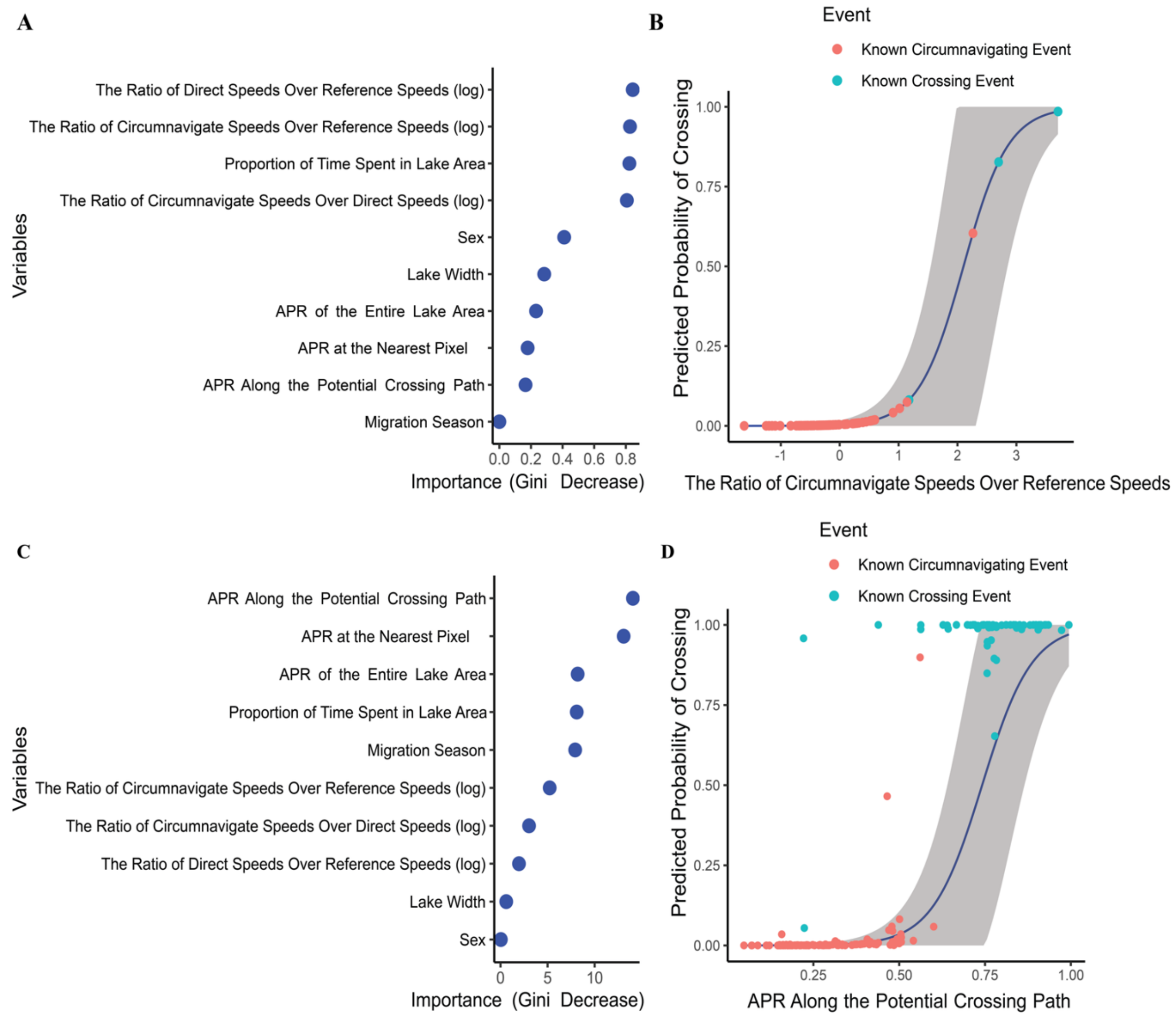


***Figure S1.*** *Model fitting results from Random Forest (RF) and Binomial Logistic (BL) models in the "Fall Only" and "Combined" datasets. All models were trained on Crossing (green) and Circumnavigating (red) events. (A) Gini Importance Index (Mean Decrease Gini Score) from the RF model classifying fall transit events. (B) BL model trained on the fall dataset*

*showing crossing probability increasing with higher log ratios of circumnavigate speed over reference speed (blue line with 95% CI in grey). (C) Gini Importance Index from the RF model trained on the combined dataset. (D) BL model trained on the combined dataset showing crossing probability increasing with APR along the potential crossing path (blue line with 95% CI in grey), with other variables fixed at median values.*

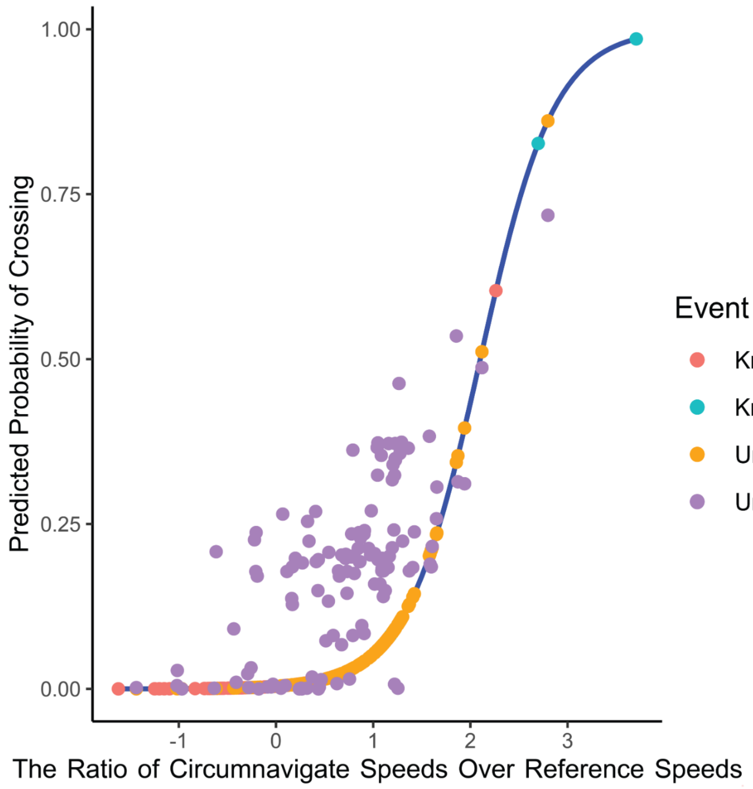


***Figure S2.*** *Relationship between crossing probability and log results of circumnavigate speeds over reference speeds in the BL model (blue line), trained on Crossing Events (green points) and Circumnavigating Events (red points). Higher ratio of speeds increases the predicted crossing probability (0 to 1). Both models (Random Forest in purple, Binomial Logistic in yellow) predict crossing probabilities for Unknown Events above 50%, demonstrating high accuracy and agreement in classifying caribou movement behaviors during fall migration.*

https://cran.r-project.org/web/packages/MuMIn/index.html.

### Dynamic Lake Ice Conditions Shape Caribou Water-Crossing Behavior in the Arctic

Qianru Liao, Eliezer Gurarie, and William F. Fagan

Appendix S6

**Appendix S6: Interannual Variability in Lake-wide Pixel- Level Ice Conditions Relative to the 56$^{th}$ Percentile Albedo Threshold**

The 56th percentile is used here as a reference quantile corresponding to the behavioral-derived threshold identified in spring migration BL models, rather than as a re-estimated or lake-scale behavioral threshold. Importantly, this percentile refers to a within-year, pixel-specific rank of albedo values, not to an absolute albedo magnitude (e.g., 0.56). At the system level, we use this reference quantile to summarize lake-wide interannual variability in (i) the absolute albedo value corresponding to the annual 56th percentile and (ii) the timing at which pixels reach this relative threshold across Contwoyto Lake.

To characterize the spring melt-season temporal dynamics of ice conditions relative to this behavioral threshold, we analyzed data from all pure-water pixels across Contwoyto Lake (N = 179, 724) over the study period (2005–2021). For each pixel and year, the threshold-crossing date, defined as the first day during the spring melt-down period when pixel-level albedo percentile rank (APR) fell below its annual 56th percentile, was identified only within the monotonic melt phase, defined as the interval from the start of the spring analysis window (DOY 92) to the annual minimum albedo, thereby avoiding confounding effects of later-season refreezing.

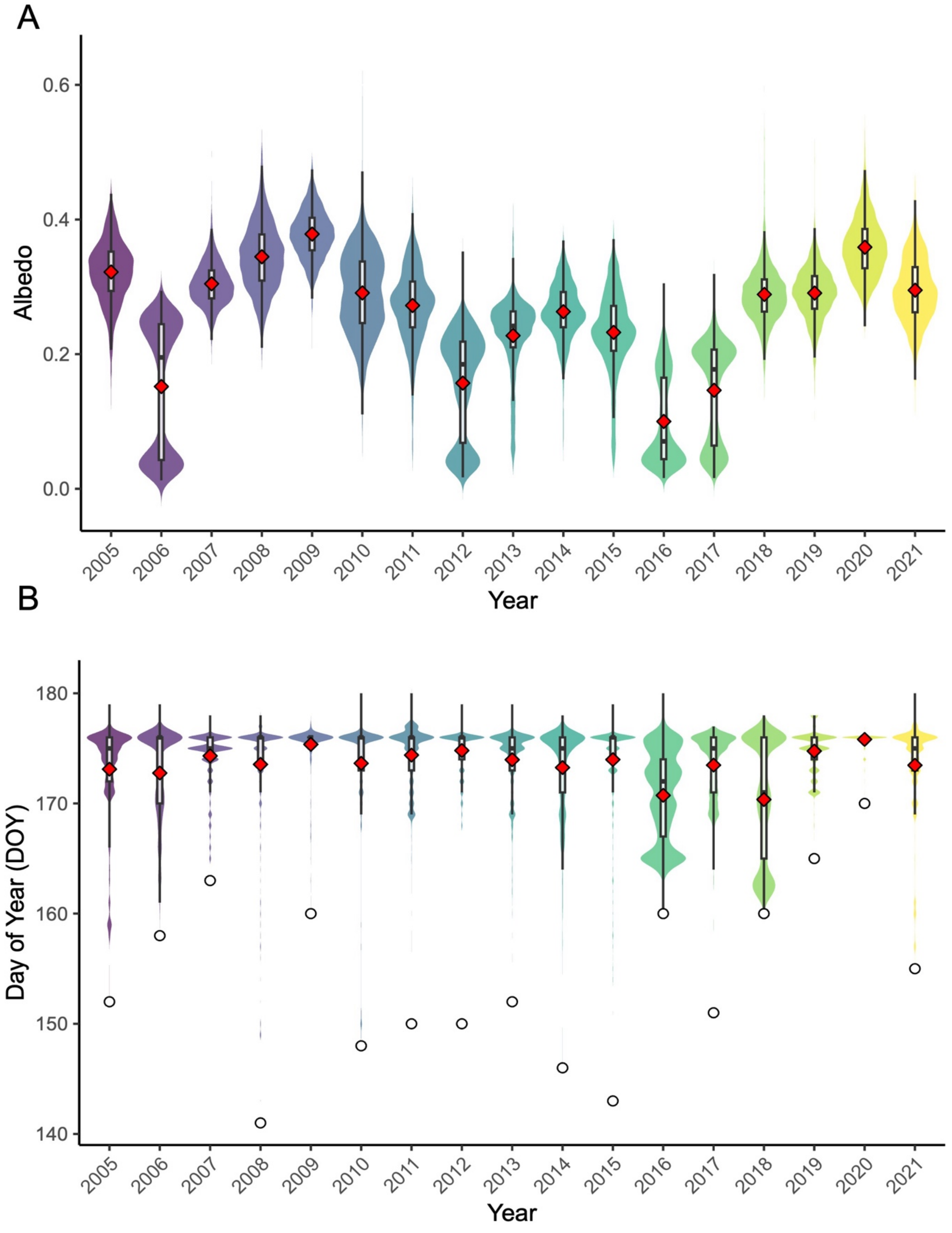


***Figure S1.*** *Lake-wide interannual variability in annual 56th-percentile albedo values and the timing of threshold crossing across Contwoyto Lake (2005–2021). (A) Lake-wide distribution of pixel-level annual 56th-percentile albedo values across Contwoyto Lake for each year, shown as violin plots with embedded boxplots; red diamonds denote yearly means. (B) Lake-*

*wide distribution of the first day of year (DOY) during the spring melt-down period when each lake pixel's albedo percentile rank (APR) first fell below its annual 56th percentile (i.e., the onset of "below-threshold" conditions), shown as violin plots with embedded boxplots; red diamonds denote yearly means, and white points indicate the earliest (minimum) DOY observed each year. The pronounced mode near DOY ~176 reflects a radiatively constrained, system-level transition in the deep central basin that occurs at a similar calendar date in most years, with only minor deviations in a small number of years. In contrast, the biologically relevant overlap with spring migration is expressed in the early tail of the distribution (e.g., as early as DOY 141, May 20 in 2008), as well as in anomalous years where the entire distribution shifts earlier (e.g., 2016).*

We fitted two types of linear models for both albedo values and the timing at which pixel-level albedo percentile rank (APR) first dropped below its annual $56^{th}$ threshold (Table S1): (1) a continuous trend model treating Year as a numeric regressor to test for linear climate warming, and (2) a categorical model treating Year as a factor to assess interannual variability.

**Table S1**. Comparison of linear trend (Year as numeric) and interannual variability (Year as factor) models for lake-wide pixel-level ice conditions (2005–2021).

| **Response Variable** | **Model** | **Predictor** | **$R^2$** | **Effect** | **P-value** |
|---|---|---|---|---|---|
| Albedo Value (56th percentile) | Linear Trend | Year (numeric) | 0.006 | -0.0015 (albedo unit /year) | < 0.001 |
| | Interannual Variability | Year (factor) | 0.647 | NA | < 0.001 |
| Albedo Threshold Timing (Day of Year, 56th percentile) | Linear Trend | Year (numeric) | 0.0005 | -0.019 (days/year) | < 0.001 |
| | Interannual Variability | Year (factor) | 0.117 | NA | < 0.001 |

Linear regression analyses revealed statistically significant but biologically negligible long-term trends, attributable to the large sample size. Annual 56th-percentile albedo values showed a slight but statistically significant decline (Slope = -0.0015/year, P < 0.001), indicating a very small long-term shift toward lower surface reflectance, consistent with marginally more ice-free or melt-advanced surface conditions. However, "Year" explained less than 1% of the total variance (($R^2$ = 0.006). Similarly, the linear trend for the timing at which pixel-level albedo percentile rank (APR) first dropped below its annual 56th percentile was statistically significant (P < 0.001) but extremely small (Slope = -0.019 day /year), corresponding to a total shift of < 0.4 days over the study period ($R^2$ = 0.0005).

In contrast, treating Year as a categorical factor substantially improved model fit, indicating that ice conditions are dominated by interannual fluctuations rather than a monotonic trend (Figure S1). The categorical model explained 65% of the variance in annual 56th - percentile albedo values ($R^2$ = 0.65, P < 0.001), with pronounced deviations from the 2005 baseline in specific years. The strongest negative albedo anomalies occurred in 2016 (β = -0.22, SE = 0.0008) and 2017 (β = -0.18, SE = 0.0008). For threshold timing, the categorical model explained 12% of the variance ($R^2$ = 0.12, P < 0.001), reflecting substantial year-to-year fluctuations relative to the weak monotonic trend. For example, 2018 exhibited significantly earlier threshold timing (β = -2.76 days), while 2020 showed significantly delayed timing (β = +2.71 days) relative to the 2005 baseline.

Overall, these results suggest that, over the 2005–2021 study period, variation in lake ice conditions surrounding the behaviorally derived albedo threshold was expressed primarily through strong interannual volatility, with little evidence for a monotonic long-term trend.

To further characterize how this variability manifests across space, we examined the spatial distributions of pixel-level annual 56th-percentile albedo values and the timing at which pixels reached the APR = 0.56 threshold across years (Figures S2 and S3).

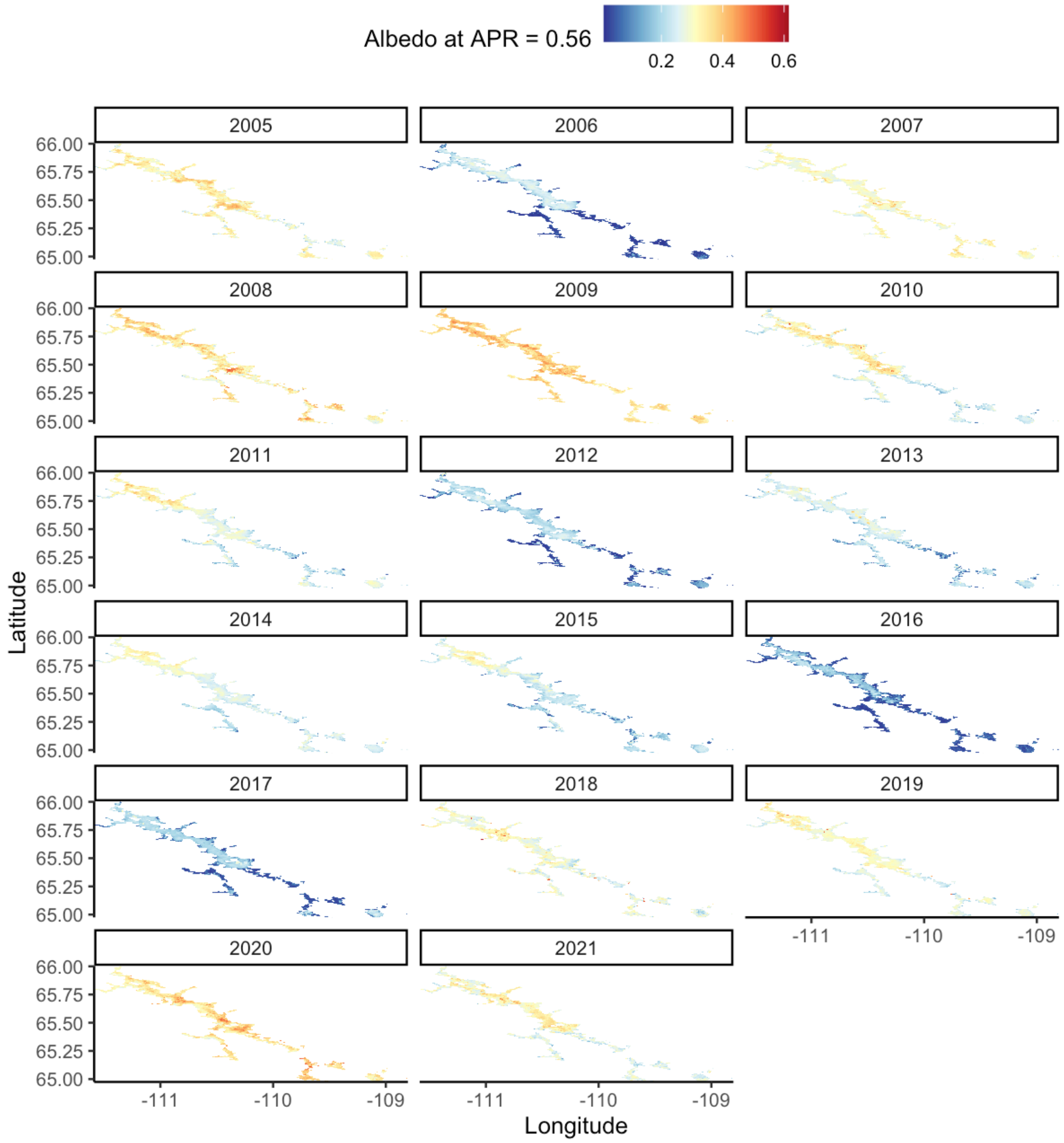


***Figure S2.*** *Spatial distribution of pixel-level annual 56th-percentile albedo values across Contwoyto Lake from 2005-2021. Colors indicate albedo values, mapped using a fixed scale from 0 (dark blue) to 1 (dark red).*

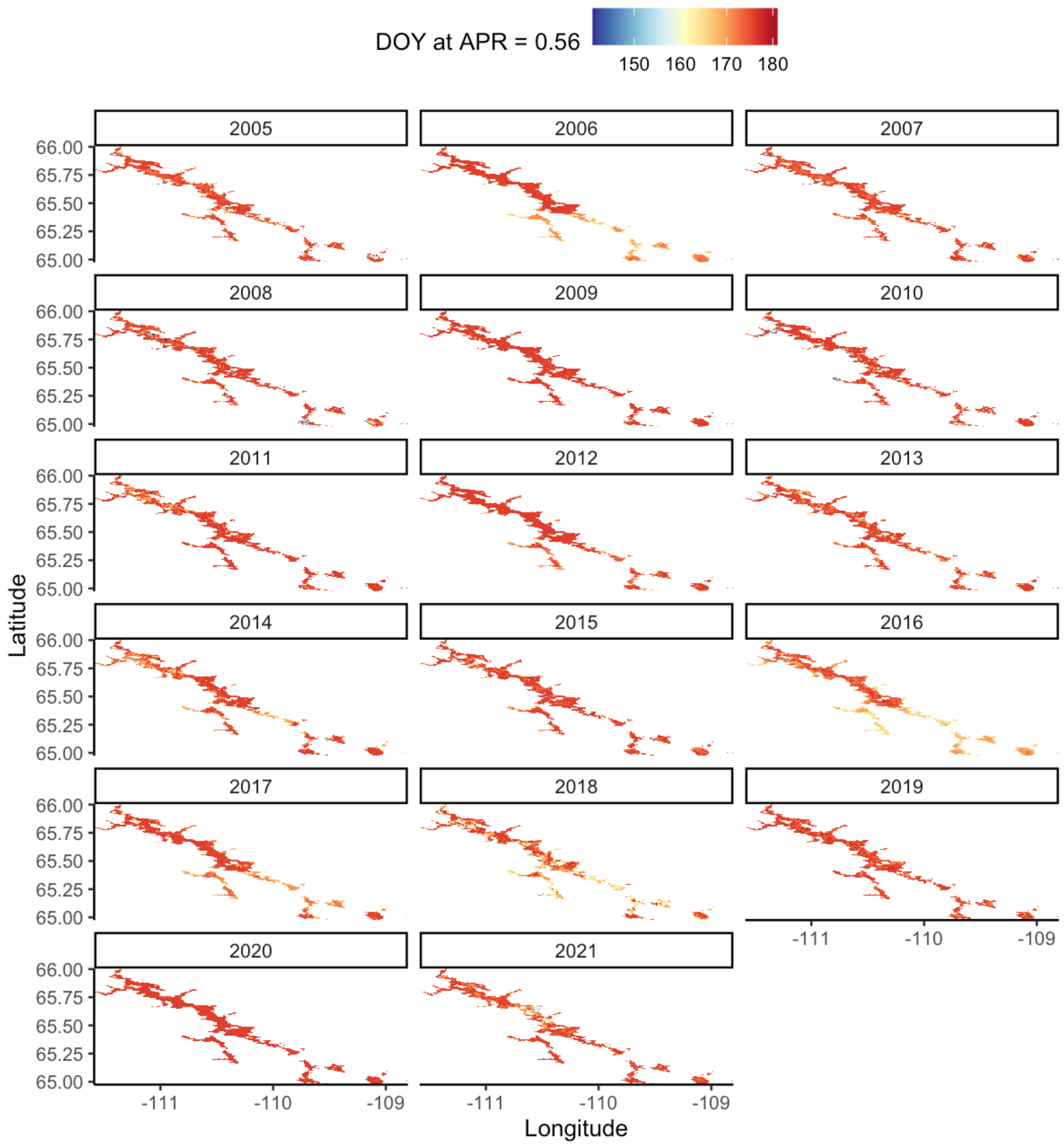


***Figure S3.*** *Spatial distribution of the first day of year (DOY) during the spring melt-down period when each lake pixel's albedo percentile rank (APR) first fell below its annual 56th percentile, indicating the onset of "below-threshold" conditions, across Contwoyto Lake from 2005–2021. Colors indicate the DOY of threshold crossing, ranging from DOY 141 (dark blue) to DOY 181 (dark red).*

Figure S2 exhibits pronounced interannual variability in the absolute albedo values associated with the annual 56th percentile. These values vary substantially among years, with some years exhibiting markedly lower albedo levels (e.g., 2006, 2016) and others showing higher values (e.g., 2009, 2020). Such variation is consistent with year-to-year differences in background surface and snow–ice conditions during spring melt, which influence the absolute albedo baseline against which relative melt progression is assessed. Because the APR-based method defines thresholds relative to each year's internal albedo distribution, Figure S2 demonstrates how the absolute albedo value corresponding to the 56th percentile varies across years with differing baseline conditions, rather than representing a single fixed physical cutoff.

In contrast, Figure S3 reveals strong temporal coherence in threshold timing across the central lake basin. Despite substantial interannual variability in absolute albedo magnitude (Figure S2), the day of year (DOY) at which pixels reach the APR = 0.56 threshold remains tightly clustered in most years. This consistency suggests that progression through the middle stages of the melt season is governed primarily by the seasonal radiative cycle, rather than by short-term meteorological variability alone.

Notable departures from this pattern occur in a small number of years (e.g., 2016 and 2018), in which a greater fraction of lake pixels reaches the threshold at earlier (lower DOY) values than in typical years. These years reflect more spatially extensive early melt conditions, during which early threshold crossing is no longer confined to a limited set of localized corridors but emerges across broader regions of the lake.

Overall, threshold timing exhibits a stable central tendency for the central lake basin in most years, reflecting radiatively constrained melt progression. Interannual variability, however, is expressed through spatial redistribution of early threshold crossing, particularly through changes in the spatial extent of early threshold crossing, with early melt affecting a larger fraction of lake pixels rather than being restricted to a few localized regions, and through pronounced deviations in anomalous years. These departures, rather than shifts in the lake-wide mean, represent the dominant source of variability most relevant to changes in the seasonal alignment between lake ice conditions and spring migration timing.
